# Early steps of the biosynthesis of the anticancer antibiotic pleurotin

**DOI:** 10.1101/2024.08.27.609176

**Authors:** Jack A. Weaver, Duha Alkhder, Panward Prasongpholchai, Michaël D. Tadesse, Emmanuel L. de los Santos, Lijiang Song, Christophe Corre, Fabrizio Alberti

**Affiliations:** School of Life Sciences, University of Warwick, Coventry, CV4 7AL, UK; Leicester Medical School, University of Leicester, Leicester, LE1 7RH, UK; UCB Biopharma, 216 Bath Road, Slough, SL1 3WE, UK; Department of Chemistry, University of Warwick, Coventry, CV4 7AL, UK

## Abstract

Pleurotin is a meroterpenoid specialised metabolite made by the fungus *Hohenbuehelia grisea*, and it is a lead anticancer molecule due to its irreversible inhibition of the thioredoxin-thioredoxin reductase system. Total synthesis of pleurotin has previously been achieved, including through a stereoselective route, however its biosynthesis has not been characterised. In this study, we used isotope-labelled precursor feeding to show that the non-terpenoid quinone ring of pleurotin and its congeners is derived from phenylalanine. We sequenced the genome of the pleurotin-producing fungus and used comparative transcriptomics to identify putative genes involved in pleurotin biosynthesis. Additionally, the heterologous expression of a UbiA-like prenyltransferase from *H. grisea* resulted in the isolation and characterisation of the first predicted pleurotin biosynthetic intermediate, 3-farnesyl-4-hydroxybenzoic acid. This work sets the foundation to fully elucidate the biosynthesis of pleurotin and its congeners, with long-term potential to optimise their production for therapeutic use and engineer the pathway towards the biosynthesis of valuable analogues.

## Introduction

Fungi are prolific producers of specialised metabolites, which are used for multiple applications including in medicine, such as β-lactam antimicrobials, cholesterol-lowering statins, and in enhancing crop yields, such as insecticide compounds made by entomopathogenic fungi.^1^ Among fungi, the mushroom-forming Basidiomycota are known to make a variety of structurally diverse compounds, including terpenoids, polyketides and amino acid-derived specialised metabolites,^2^ some of which have high potential to be developed into drugs. However, this process is often hampered because of inherent difficulties connected with slow growth and challenging genetic engineering of Basidiomycota fungi.^3^ Understanding the biosynthesis of bioactive natural products from fungi of this division can allow us to direct it to the production of valuable analogues and congeners, as seen in the case of the pleuromutilin antibiotics.^4,5^

Among Basidiomycota fungi, the pleurotoid mushroom *Hohenbuehelia grisea* makes the anticancer antibiotic pleurotin, a meroterpenoid natural product that was first discovered in 1947 and shown to inhibit the growth of *Staphylococcus aureus*.^6^ Pleurotin was later proven to inhibit the growth of some fungi^7,8^ and to have antitumor activity through potent irreversible inhibition of the thioredoxin-thioredoxin reductase system, which makes it a lead anticancer compound.^9^

Considerable work has emerged in the past years on the study of bioactive pleurotin congeners made by *H. grisea* (Figure 1) including the study of the antiviral 4-hydroxypleurogrisein^10^ and the isolation of cysteine-derived analogues of pleurotin.^11^ An optimised fermentation process for the production of pleurotin in high titres (more than 300 mg l^-1^) has been developed.^12^ The total synthesis of (±)-pleurotin has been achieved through 26^13^ and 13 linear steps,^14^ as well as of its congeners (±)-pleurogrisein and (±)-4-hydroxypleurogrisein.^15^ Importantly, the stereoselective syntheses of (−)-pleurotin and its congeners (+)-leucopleurotin, (+)-dihydropleurotinic acid, and (+)-leucopleurotinic acid were recently reported in 15-16 steps,^16^ paving the way for the synthesis of new stereoselective pleurotin analogues.

**Figure 1.**
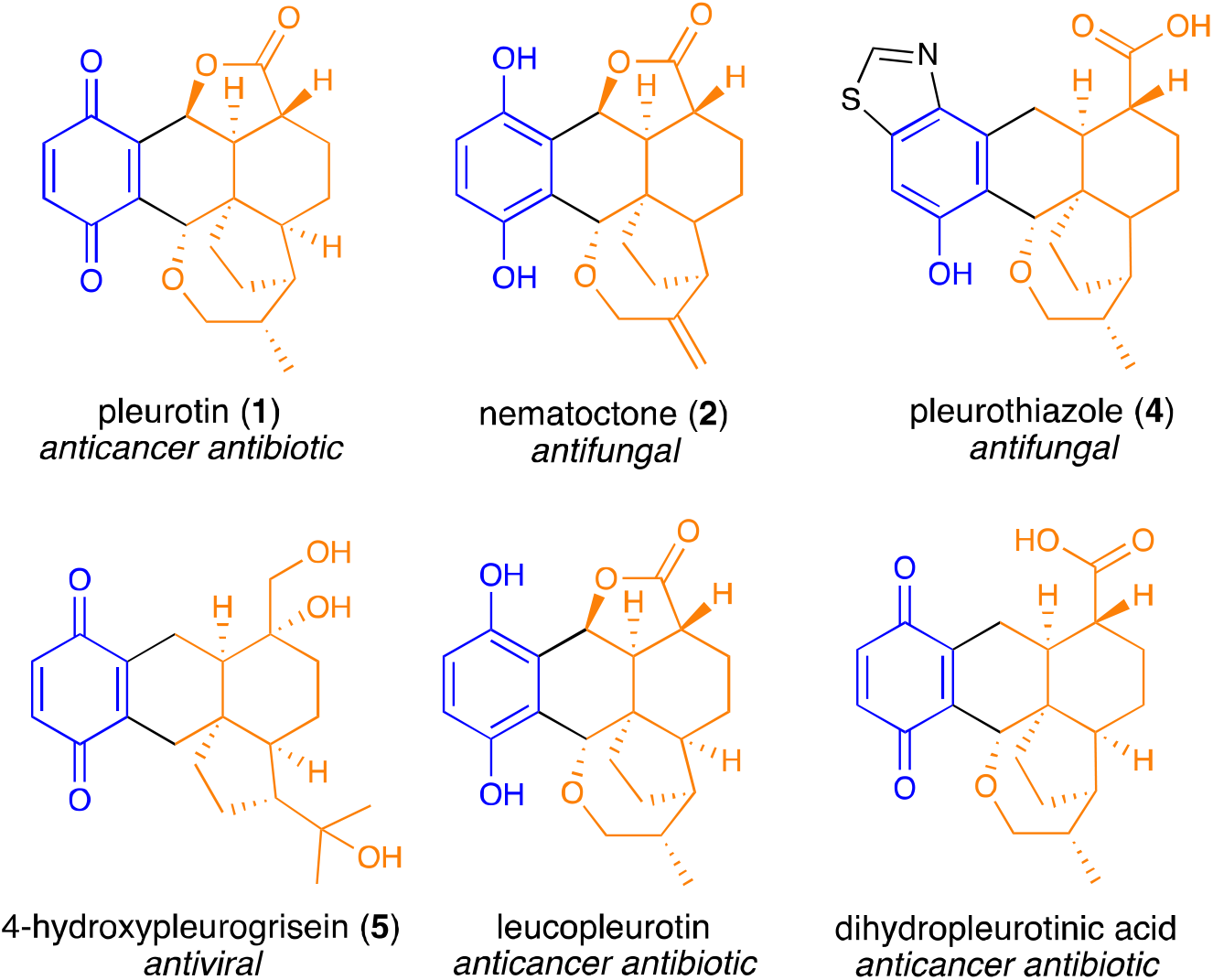
Structures of pleurotin and selected bioactive analogues. The terpenoid moiety is highlighted in orange, the non-terpenoid moiety in blue, the connecting bonds and additional groups in black.

Preliminary studies on the biosynthesis of pleurotin were conducted by Arigoni’s group through feeding of the fungus with isotope-labelled predicted precursors and intermediates, suggesting that pleurotin may derive from farnesylhydroquinone through several steps of cyclisation, rearrangement, and oxidations.^17–19^ The exact sequence of reactions that lead to pleurotin is still unknown, and so are the enzymes involved in the biosynthetic pathway.

In this study, we set out to investigate the origin of the quinone ring of pleurotin by means of isotope-labelled precursor feeding. We also sequenced the genome of the pleurotin-producing fungus *H. grisea* and performed comparative transcriptomics analysis to pinpoint candidate biosynthetic genes. Characterisation of a UbiA-like prenyltransferase (PTase) led us to isolate the first predicted intermediate in pleurotin biosynthesis. This work sets the foundation to characterise the biosynthetic pathway to pleurotin and its valuable congeners.

## Results and Discussion

### Isolation and structural characterisation of pleurotin

Firstly, we cultured *Hohenbuehelia grisea* (strain ATCC 60515) in YM glucose to confirm production of pleurotin (1) from this strain. Metabolite extracts were analysed through liquid chromatography-high-resolution mass spectrometry (LC-HRMS), in which extracted ion chromatogram at *m*/z 355.1545 (calculated for [M + H]^+^, where M= C_21_ H_22_ O_5_) showed a main species at retention time 20.5’ and a minor one at retention time 18.2’ (Supplementary Figure 1). Based on the literature, we hypothesised that the minor product is likely to be pleurotin’s structural isomer nematoctone (**2**), which is reported to be produced in sub-milligram per litre level by pleurotin-producing fungi.^10^ In order to unequivocally confirm the identity of pleurotin, we scaled up cultures of *H. grisea* in YM glucose and used a combination of flash chromatography and HPLC to purify **1**. Structural characterisation was achieved through ^1^H-NMR spectroscopy (Supplementary Figure 2), which was in agreement with literature data reported for **1**.^16^

### Biosynthetic origin of the quinone ring of pleurotin and its congeners

Meroterpenoids are hybrid natural products that include a terpenoid moiety and a non-terpenoid portion (see Figure 1). The non-terpenoid side of fungal meroterpenoids often includes a polyketide moiety, e.g. 5-methylorsellinic acid in mycophenolic acid,^20^ which is made by an iterative type-I polyketide synthase (PKS). However, fungal meroterpenoid biosynthetic pathways can sometimes use unusual starter units, such as nicotinyl-CoA in pyripyropene A biosynthesis.^21^ They can also be made independently of PKSs and use 4-hydroxybenzoic acid (4-HBA) as a precursor for their non-terpenoid moiety. For instance, the biosynthesis of antroquinonol^22^ and vibralactone^23^ involves the use of 4-HBA that can either be made through the endogenous shikimate pathway *via* chorismate or through exogenous phenylalanine.

Arigoni’s group performed preliminary studies on pleurotin biosynthesis through incorporation of [1-^13^C]- and [1,2-^13^C_2_]-acetate to show that **1** includes a C_15_ terpenoid side that derives from the mevalonate pathway, whereas its quinone ring was shown to derive from 4-hydroxybenzoic acid (4-HBA) through incorporation of a deuterated analogue of it.^18,19^ In order to shed light on the biosynthetic origin of the non-terpenoid side of pleurotin, we decided to test whether phenylalanine **(3)** can provide the quinone ring of **1**. We fed cultures of *H. grisea* with *L*-phenyl-^13^C_9_-alanine and analysed ethyl acetate crude metabolite extracts through LC-HRMS, using *H. grisea* grown in standard *L*-phenylalanine as a control (Figure 2). The extract of *H. grisea* fed with *L*-phenyl-^13^C_9_-alanine showed a species with *m*/z 361.1730 (calculated *m*/z of 361.1746) at retention time 20.5’ (Figure 2C) corresponding to a six-Da increase compared to the pleurotin peak with *m*/z 355.1530 (calculated *m*/z of 355.1545), and absent in *H. grisea* grown in standard *L*-phenylalanine (Figure 2B). MS^2^ spectra were analysed to further confirm the incorporation of the six heavy carbons in **1** (Supplementary Figure 3). We also investigated the incorporation of the heavy carbons provided by *L*-phenyl-^13^C_9_-alanine in other pleurotin-related metabolites that we could observe within the extract. We were able to detect the same six-Da shift for the additional species with m/z 355.1530 at retention time 18.2’ (Supplementary Figure 4), a species with *m*/z 400.1568 at retention time 19.4’ (Supplementary Figure 5), and one with *m*/z 361.1996 at retention time 16.3’ (Supplementary Figure 6), which returned predicted molecular formulae corresponding to those of the known pleurotin congeners nematoctone **(2)**,^10^ pleurothiazole **(4)**,^11^ and 4-hydroxypleurogrisein **(5)**,^10^ respectively. High percentages of ^13^C incorporation from *L*-phenyl-^13^C_9_-alanine into **1, 2, 4** and **5** were detected, ranging from 63 to 68% compared to the corresponding ^12^C-containing species (Supplementary Table 1).

**Figure 2.**
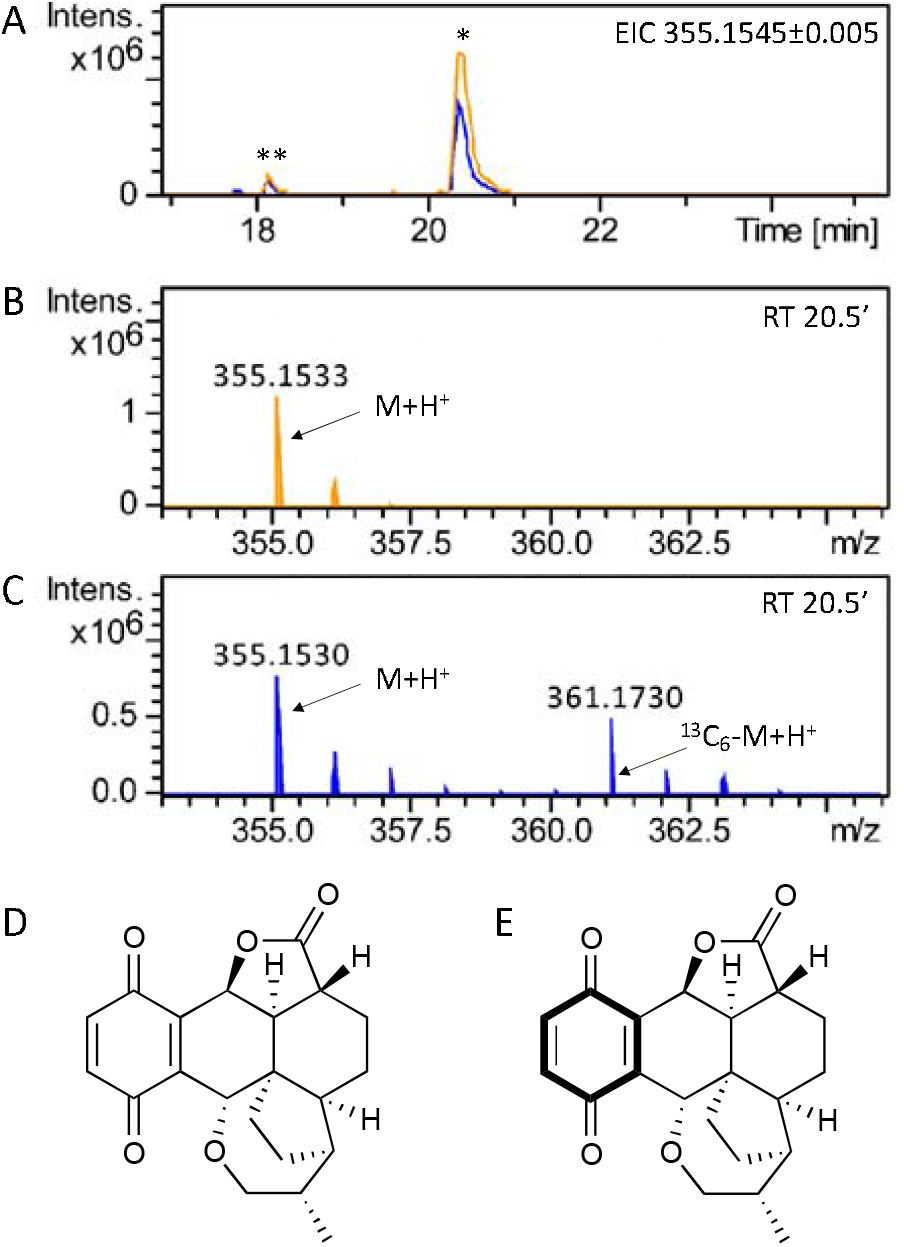
LC-HRMS detection of pleurotin **(1)** from *H. grisea* fungal extract. **A)** Extracted ion chromatogram in positive mode at *m/z* = 355.1545±0.005 is shown, highlighting accumulation of pleurotin in *H. grisea* crude extracts grown in standard L-phenylalanine (trace in orange) and in *L*-phenyl-^13^C_9_-alanine (trace in blue). One major peak (*) at retention time 20.5’ is seen, corresponding to pleurotin **(1)**, based on NMR characterisation of the purified compound; one additional peak (**) at retention time 18.2’ is seen, likely corresponding to nematoctone **(2)**.^10^ **B)** Mass spectrum of pleurotin (M= C_21_H_22_O_5_) detected at retention time 20.5’; *m/z* calculated for [M + H]^+^= 355.1545. **C)** Mass spectrum of pleurotin and ^13^C_6_-pleurotin detected at retention time 20.5’; *m/z* calculated for [^13^C_6_-M + H]^+^= 361.1746. **D)** Pleurotin. **E)** ^13^C_6_-pleurotin.

Based on stable isotope feeding, we propose that *L*-phenylalanine (**3**) can provide the quinone ring of **1** and of its congeners **2, 4** and **5** (Scheme 1). A phenylalanine ammonia lyase, which is sometimes found in fungal meroterpenoid BGCs,^24,25^ is likely to convert **3** into trans-cinnamic acid, which can then undergo hydroxylation to make 4-coumaric acid, followed by side-chain degradation to give 4-HBA. We then predict the farnesylation of 4-HBA to be catalysed by a UbiA-like PTase, a widely characterised enzyme class involved in the biosynthesis of fungal meroterpenoids such as mycophenolic acid,^20^ anditomin,^26^ ascochlorin and ascofuranone,^27^ to name a few.

### Whole-genome sequencing of *Hohenbuehelia grisea*

Since no genome sequence was publicly available for pleurotin-producing fungi, we next aimed to sequence the genome of *H. grisea* ATCC 60515, so that it would serve as a framework to look for the biosynthetic genes involved in the production of pleurotin. Genomic DNA from *H. grisea* was purified and subjected to whole-genome sequencing using a combination of nanopore long-read sequencing and Illumina short-read sequencing, as well as Illumina-sequenced mRNA for improved gene annotation for Funannotate training.^28^ The assembled genome was deposited on NCBI under accession number JASNQZ000000000, BioProject: PRJNA956249. The genome had a total length of 38.87 Mb and comprised of 25 contigs (N_50_ 2.88 Mb). Full assembly statistics are reported in Supplementary Table 2. The completeness of the genome assembly and annotation were assessed using BUSCO, which returned a high score of 97.3% for the genomic scaffold, and of 94.3% for the predicted proteome (see full BUSCO results in Supplementary Table 3). A total of 14,934 genes were predicted to be in the genome, out of which 14,602 were predicted to be protein-coding. On average, each gene was anticipated to include 6.72 exons. Additionally, 332 tRNA genes were predicted.

### Analysis of *Hohenbuehelia grisea* secondary metabolite biosynthetic gene clusters

The biosynthetic enzymes that produce fungal meroterpenoids are often encoded by genes that are co-localised in biosynthetic gene clusters (BGCs).^29^ We therefore analysed the assembled and annotated genome of *H. grisea* for the presence of secondary metabolite gene clusters using fungiSMASH,^30^ which returned 21 predicted BGCs (see Table 1). The biosynthesis of meroterpenoid natural products like **1** generally involves signature biosynthetic genes such as the aforementioned UbiA-like PTase for prenylation of a non-terpenoid moiety, as well as a transmembrane terpene cyclase (TC) for cyclisation. A PKS can also be present in the BGC if the meroterpenoid includes a polyketide non-terpenoid moiety.^29^

**Table 1.**
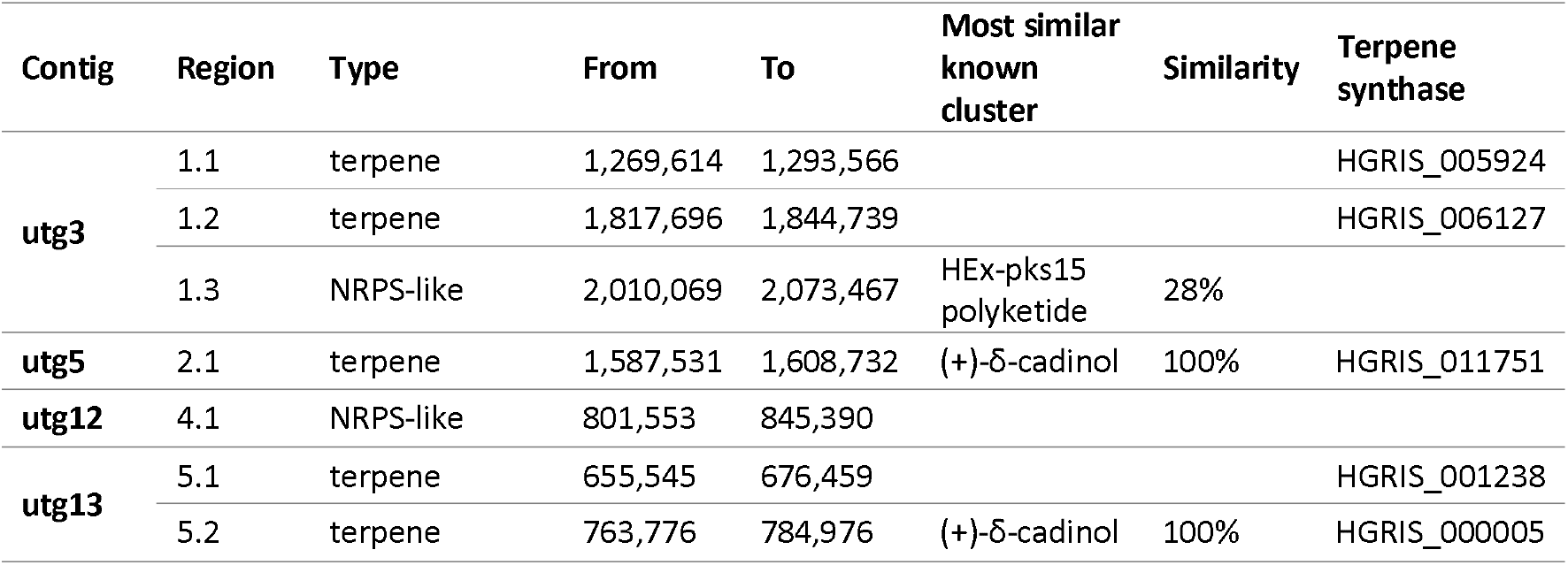
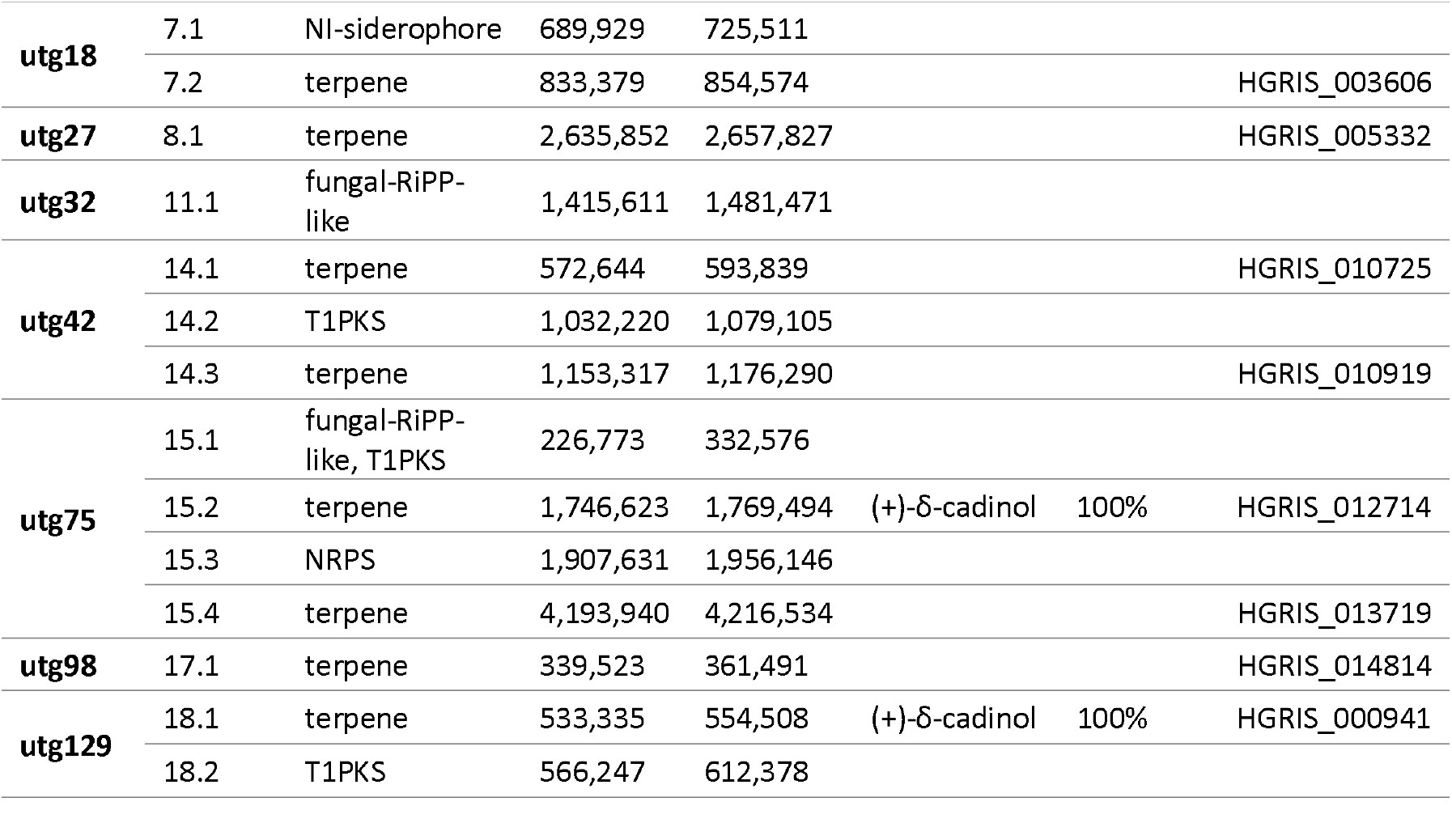
Secondary metabolite analysis of the *H. grisea* genome performed through FungiSMASH 7.0. NRPS= non-ribosomal peptide synthetase; NI-siderophore= NRPS-independent, IucA/IucC-like siderophore; RiPP= ribosomally synthesised and post-translationally modified peptide; T1PKS= type I polyketide synthase.

| Contig | Region | Type | From | To | Most similar known cluster | Similarity | Terpene synthase |
| --- | --- | --- | --- | --- | --- | --- | --- |
| utg3 | 1.1 | terpene | 1,269,614 | 1,293,566 |  |  | HGRIS_005924 |
|  | 1.2 | terpene | 1,817,696 | 1,844,739 |  |  | HGRIS_006127 |
|  | 1.3 | NRPS-like | 2,010,069 | 2,073,467 | HEx-pks15 polyketide | 28% |  |
| utg5 | 2.1 | terpene | 1,587,531 | 1,608,732 | (+)-δ-cadinol | 100% | HGRIS_011751 |
| utg12 | 4.1 | NRPS-like | 801,553 | 845,390 |  |  |  |
| utg13 | 5.1 | terpene | 655,545 | 676,459 |  |  | HGRIS_001238 |
| | 5.2 | terpene | 763,776 | 784,976 | (+)-δ-cadinol | 100% | HGRIS_000005 |
| **utg18** | 7.1 | Nl-siderophore | 689,929 | 725,511 |  |  |  |
|  | 7.2 | terpene | 833,379 | 854,574 |  |  | HGRIS_003606 |
| **utg27** | 8.1 | terpene | 2,635,852 | 2,657,827 |  |  | HGRIS_005332 |
| **utg32** | 11.1 | fungal-RiPP-like | 1,415,611 | 1,481,471 |  |  |  |
| **utg42** | 14.1 | terpene | 572,644 | 593,839 |  |  | HGRIS_010725 |
|  | 14.2 | T1PKS | 1,032,220 | 1,079,105 |  |  |  |
|  | 14.3 | terpene | 1,153,317 | 1,176,290 |  |  | HGRIS_010919 |
| **utg75** | 15.1 | fungal-RiPP-like, T1PKS | 226,773 | 332,576 |  |  |  |
| | 15.2 | terpene | 1,746,623 | 1,769,494 | (+)-δ-cadinol | 100% | HGRIS_012714 |
|  | 15.3 | NRPS | 1,907,631 | 1,956,146 |  |  |  |
|  | 15.4 | terpene | 4,193,940 | 4,216,534 |  |  | HGRIS_013719 |
| **utg98** | 17.1 | terpene | 339,523 | 361,491 |  |  | HGRIS_014814 |
| **utg129** | 18.1 | terpene | 533,335 | 554,508 | (+)-δ-cadinol | 100% | HGRIS_000941 |
|  | 18.2 | T1PKS | 566,247 | 612,378 |  |  |  |

Upon close inspection of the BGCs detected through fungiSMASH in the genome of *H. grisea*, including the 12 terpene BGCs, no UbiA-like PTase gene could be identified. We therefore searched for UbiA-like PTases within the genome of *H. grisea* through local BlastP against the predicted proteome of the fungus, using the sequence of the *Saccharomyces cerevisiae* UbiA-like PTase COQ2 (para hydroxybenzoate:polyprenyltransferase)^31^ as the query. We found four high-scoring homologous predicted proteins to COQ2 in *H. grisea*, HGRIS_009700, HGRIS_000873, HGRIS_000139 and HGRIS_006930, which share homology with COQ2 of 45%, 33%, 33% and 36%, respectively (see Supplementary Table 4). Similarly, we searched for homologues of the FAD-binding mono-oxygenase VibMO1, which is responsible for the oxidative decarboxylation of prenyl 4-HBA as part of vibralactone biosynthesis in the Basidiomycota fungus *Boreostereum vibrans*.^32^ An enzyme homologous to VibMO1 can be predicted to decarboxylate 3-farnesyl-4-HBA into farnesylhydroquinone in the early steps of pleurotin biosynthesis (Figure 2). Seven high-scoring homologous proteins to VibMO1 were found in *H. grisea*, HGRIS_002929, HGRIS_001008, HGRIS_001007, HGRIS_004802, HGRIS_014061, HGRIS_008360 and HGRIS_010717 (Supplementary Table 5). None of these appeared to be located in the putative terpenoid BGCs detected through FungiSMASH.

### Comparative transcriptomics analysis enables the shortlisting of putative pleurotin biosynthesis genes

In order to investigate the involvement of the putative biosynthetic genes found through FungiSMASH and local BlastP in the biosynthesis of **1**, we performed comparative transcriptomics analysis in *H. grisea* cultures that exhibited differential pleurotin production. Specifically, cultures of *H. grisea* were grown in shake-flask in media containing five different carbon sources (glucose, mannitol, fructose, galactose and lactose - based on Robbins *et al*.^6^) in parallel. Ethyl acetate metabolite extracts arising from 2-week-old fungal cultures were analysed using LC-HRMS, showing that only cultures grown in glucose as the carbon source were able to produce **1** in appreciable amounts, which was readily detected as a peak with *m/z* 355.1532 at retention time 20.5’ (Supplementary Figure 7). Fungi grown in fructose as the carbon source showed limited production of pleurotin, whereas cultures grown in mannitol, fructose and galactose did not produce pleurotin at all.

Once differential production of **1** could be ascertained in different culturing media, glucose and mannitol were picked as the two carbon sources to perform RNAseq analysis at two timepoints, seven and 14 days, based on the observation that accumulation of **1** could be detected starting from as early as one-week when culturing *H. grisea* in YM glucose. On day seven, 512 genes were overexpressed in glucose-grown mycelia compared to mannitol-grown mycelia (3.43% of the total number of genes) (Supplementary Dataset 1). At day 14, the number of overexpressed genes between glucose-grown and mannitol-grown mycelia had increased to 948 (6.35% of the total number of genes). The expression of the *H. grisea* UbiA-like PTase homologue genes was examined, which showed that only *HGRIS_000139* was upregulated during production of **1** at seven days (showing a Log_2_FC of 1.68), whereas the three other PTases were downregulated in glucose-grown compared to mannitol-grown fungi at both timepoints (Supplementary Figure 8). Looking at the *H. grisea* homologues of FAD-binding mono-oxygenase VibMO1, only gene *HGRIS_001007* appeared to be upregulated in glucose-grown compared to mannitol-grown fungi at day seven, with a Log_2_FC of 2.20 (Supplementary Figure 9).

Interestingly, most terpene synthases identified to be within *H. grisea* BGCs from FungiSMASH were downregulated when pleurotin was produced (Supplementary Figure 10), with the only exception of HGRIS_005332, which was upregulated at day 14 with a Log_2_FC of 1.54. However, BlastP analysis of the clustered genes at region 8.1 (Supplementary Table 6) excluded any involvement of nearby genes to HGRIS_005332 in the biosynthesis of secondary metabolites.

### The UbiA-like PTase HGRIS_000139 catalyses farnesylation of 4-HBA to make 3-farnesyl-4-HBA

Since the putative UbiA-like PTase *HGRIS_000139* was upregulated during pleurotin production, we aimed to investigate its function by expressing it heterologously in *Aspergillus oryzae* NSAR1.^33^ We also decided to include a copy of the *H. grisea* farnesyl pyrophosphate synthase (FPPS) gene *HGRIS_005317*, to provide increased amounts of FPP for suitable precursor supply. To this aim, we amplified the intron-less sequences of both *HGRIS_000139* and *HGRIS_005317* from the cDNA of *H. grisea* (Supplementary Figure 11) and used them to assemble an expression vector based on the pTYGSarg plasmid backbone,^34^ named pDA001 (Supplementary Figure 12). We transformed pDA001 into *A. oryzae* NSAR1, and confirmed heterologous gene insertion in *A. oryzae* DA1 through PCR amplification of HGRIS_005317 and HGRIS_000139 (Supplementary Figures 13, 14). Metabolite analysis was performed through LC-HRMS to compare the crude extracts of *A. oryzae* DA1 and *A. oryzae* NSAR1, which highlighted the accumulation of a species with *m/z* 343.2269 in *A. oryzae* DA1, absent in the recipient strain *A. oryzae* NSAR1 (Figure 3). Prediction of the most likely molecular formula suggested a species with formula C_22_H_30_O_3_, (calculated for M[C_22_ H_30_ O_3_]^+^= 343.2273), which was consistent with the formula of 3-farnesyl-4-hydroxybenzoic acid (3-farnesyl-4-HBA) **(6)**. Purification of **6** was achieved using a combination of flash chromatography and HPLC, and its structure was confirmed through ^1^H-NMR spectroscopy (Supplementary Figure 15), which was in agreement with literature data reported for **6**.^24,35^ We therefore propose that HGRIS_000139 is a UbiA-like PTase that catalyses the farnesylation of 4-HBA in *H. grisea* to give the first predicted pleurotin intermediate **6**.

**Figure 3.**
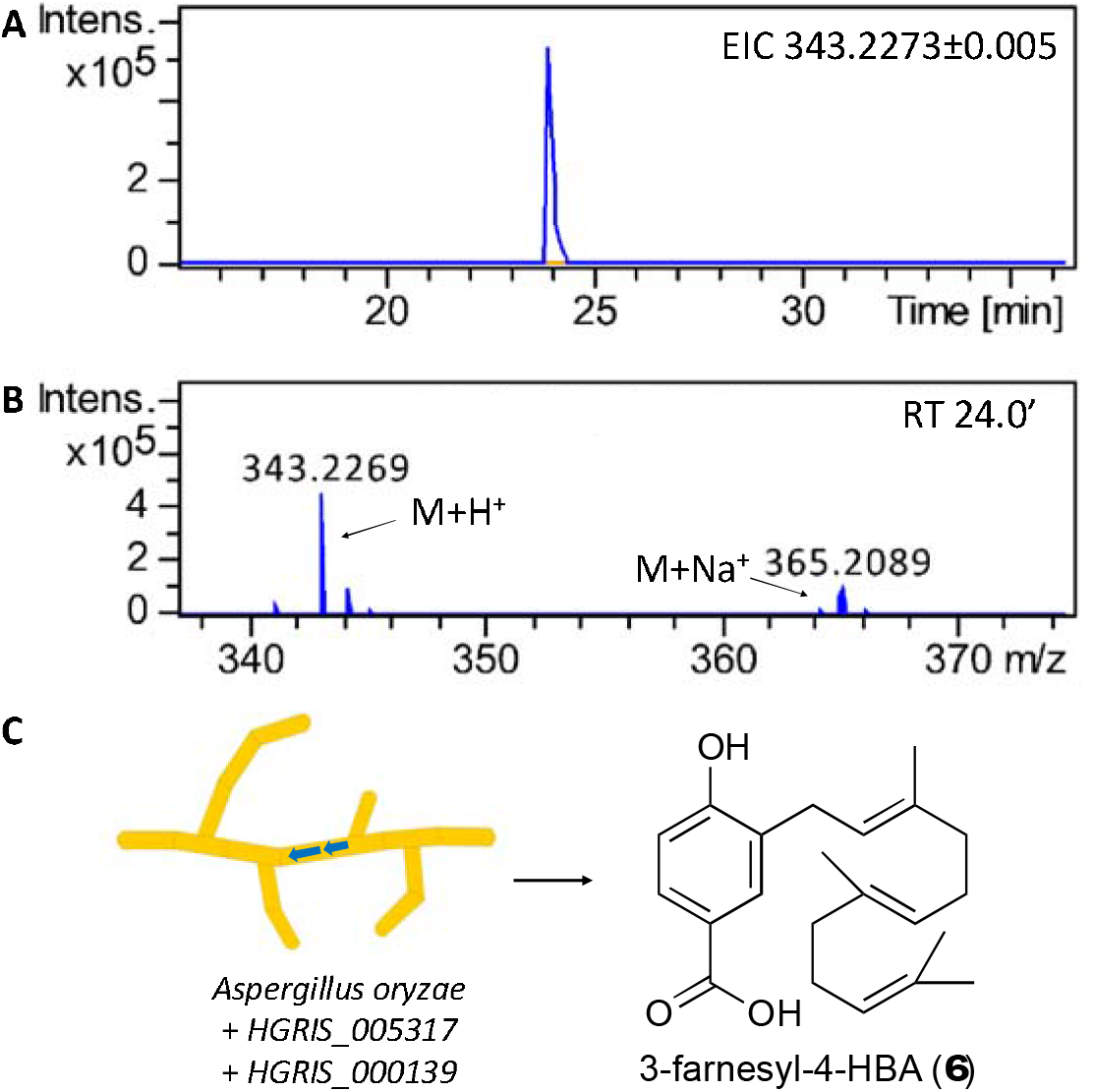
LC-HRMS detection of 3-farnesyl-4-HBA **(6)** in A. oryzae DA1. **A)** Extracted ion chromatogram in positive mode at *m/z* = 343.2273±0.005 is shown, highlighting accumulation of **6** in *A. oryzae* DA1 crude metabolite extracts (trace in blue) and its absence in *A. oryzae* NSAR1 (trace in orange). **B)** Mass spectrum of **6** (M= C_22_H_30_O_3_) detected at retention time 24.0’. C) Schematic representation of the heterologous expression of *HGRIS_005317* and *HGRIS_000139* in A. oryzae leading to the accumulation of **6**.

## Conclusions

In summary, we characterised the early steps of pleurotin biosynthesis, showing that L-phenylalanine can provide the quinone ring of **1**, likely *via* 4-HBA, and that the farnesylation of 4-HBA catalysed by the UbiA-like aPTase HGRIS_000139 leads to the first predicted pathway intermediate 3-farnesyl-4-HBA. We also sequenced the genome of the pleurotin-producing fungus *H. grisea*, providing a scaffold for the identification of the other pleurotin biosynthetic genes. FungiSMASH detection of putative BGCs did not conclusively point toward a candidate pleurotin BGC. Examples are known of fungal meroterpenoids that are produced by enzymes that are not colocalised, such as vibralactone,^36^ or are made by enzymes encoded by genes that are split across two BGCs, such as ascofuranone.^27^ Comparative transcriptomics analysis allowed us to generate a list of candidate biosynthetic genes that can be further characterised to determine their involvement in pleurotin biosynthesis and obtain a full picture of the pathway.

**Scheme 1.**
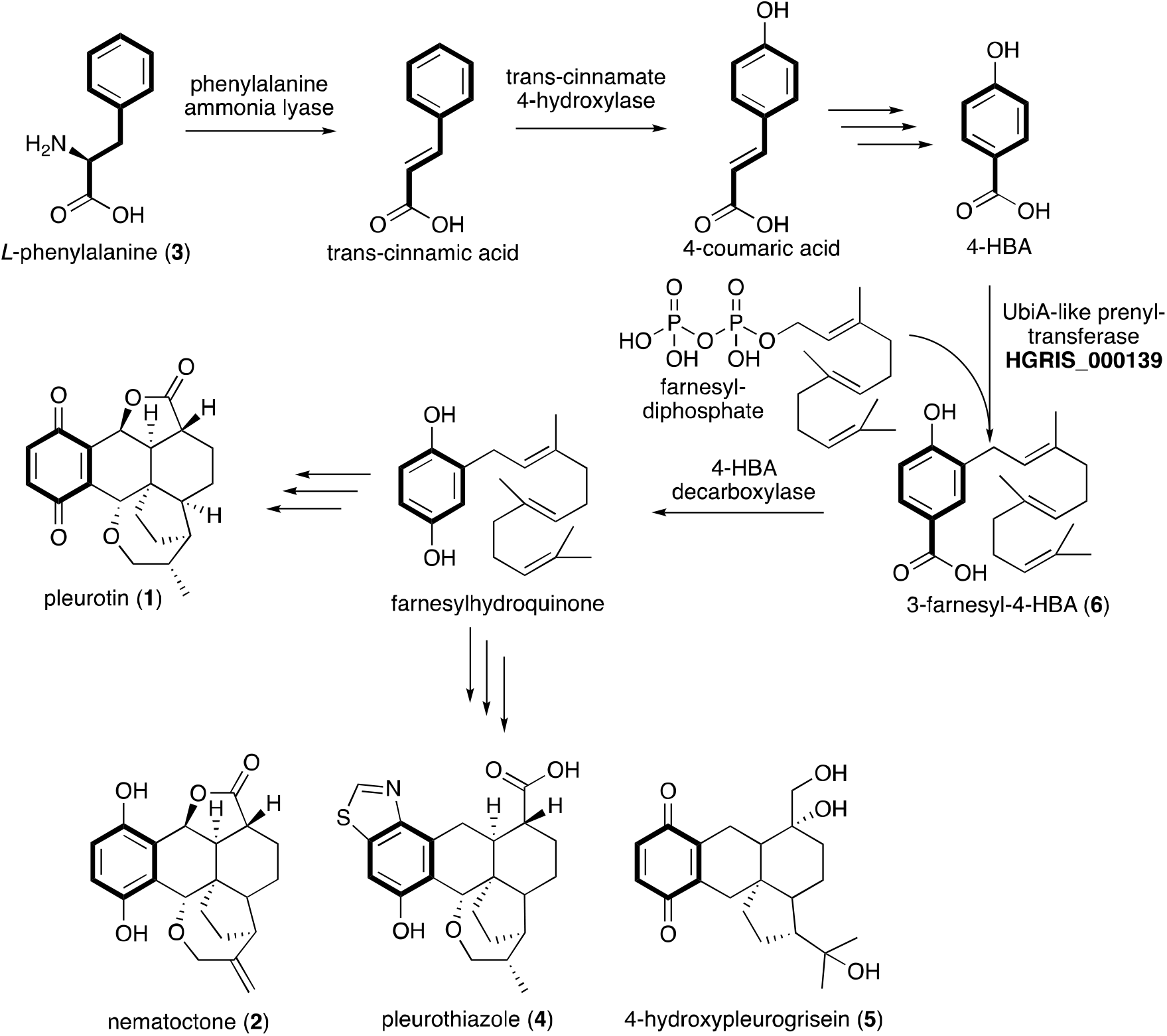
Proposed biosynthesis of pleurotin and selected congeners in *H. grisea*.

## Author contributions

J.W., D.A., P.P. and M.T. carried out experimental work. E.L.d.l.S. provided guidance on the genome assembly and annotation. L.S. performed NMR spectroscopy assignment and assisted in MS data interpretation. C.C. provided insights on the experimental design, as well as resources. F.A. conceived the study and wrote the article, with contributions from all authors.

## Data availability

The data supporting this article have been included as part of the Supplementary Information and Supplementary Dataset 1. The assembled genome was deposited on NCBI under accession number JASNQZ000000000, BioProject: PRJNA956249.

## Conflicts of interest

There are no conflicts to declare.

## Acknowledgments

J.W. was supported by a scholarship from the Engineering and Physical Sciences Research Council and the Biotechnology and Biological Sciences Research Council [EP/L016494/1] through the Centre for Doctoral Training in Synthetic Biology (SynBioCDT). M.T. was supported by a scholarship from the Midlands Integrative Biosciences Training Partnership [BB/T00746X/1], a BBSRC-funded Doctoral Training Partnership. F.A. was supported by a Leverhulme Trust Early Career Fellowship [ECF-2018-691] and a UKRI Future Leaders Fellowship [MR/V022334/1]. The authors acknowledge use of chromatography equipment provided by the Bio-Analytical Shared Resource Laboratories within the School of Life Sciences, University of Warwick. WISB, BBSRC/EPSRC Synthetic Biology Research Centre (grant ref: BB/M017982/1), is also acknowledged. Christopher de Wolf is thanked for assistance in data acquisition on chromatography equipment. Dr Marlene Rothe is thanked for assistance in NMR spectroscopy data acquisition.

## Notes

### Competing Interest Statement

The authors have declared no competing interest.

