## Supplementary information for "Early steps of the biosynthesis of the anticancer antibiotic pleurotin"

### Materials and methods

**Reagents**

Chemicals and ingredients for media used in this work were obtained from Merck, Fisher, Oxoid, ForMedium, Thermo Fisher Scientific unless otherwise stated. Media were prepared and sterilised by autoclaving using a programme at 121°C for 20 minutes. Deionised water was used for all solutions unless stated.

**Growth conditions for fungal and bacterial strains**

*Hohenbuehelia grisea* (strain ATCC 60515, also known as strain T-177) was maintained on YM glucose agar plates (3 g l^-1^ yeast extract, 3 g l^-1^ malt extract, 10 g l^-1^ D-glucose, 5 g l^-1^ bacto peptone, 20 g l^-1^ agar, pH 6.2) at 25°C.

*Aspergillus oryzae* NSAR1 (genotype niaD-, sC-, ΔargB, adeA-) was maintained on MEA plates with appropriate supplements (15 g l^-1^ malt extract, 1.5 g l^-1^ arginine, 1.5 g l^-1^ methionine, 0.1 g l^-1^ adenine, 2 g l^-1^ ammonium sulphate, 20 g l^-1^ agar) at 28°C. Propagation of plasmids was performed in *E. coli* One Shot® ccdB Survival^TM^ 2 T1R competent cells (Life Technologies), which were grown in LB agar plates (10 g l^-1^ NaCl, 10 g l^-1^ tryptone, 5 g l^-1^ yeast extract, 20 g l^-1^ agar, pH 7) with appropriate antibiotic as needed (ampicillin 100 mg l^-1^ when containing plasmids pTYGSarg and pDA001) at 37°C.

**Metabolite purification from *H. grisea* and *A. oryzae* strains**

To purify **1**, *H. grisea* was cultured in 1 l of YM glucose broth (3 g l^-1^ yeast extract, 3 g l^-1^ malt extract, 10 g l^-1^ D-glucose, 5 g l^-1^ peptone, pH 6.2) for two weeks, in an orbital shaker at 180 rpm at 25°C. To purify **5**, *A. oryzae* DA1 was grown in 1 l of CMP medium (35 g l^−1^ Czapek-dox broth, 20 g l^−1^ maltose, 10 g l^−1^ peptone) for ten days, in an orbital shaker at 180 rpm at 28°C. Metabolite extractions were performed twice using two volumes of ethyl acetate, followed by concentration of the crude extract *in vacuo*. Flash chromatography was used to fractionate the crude extract on a Biotage Selekt automated flash purification system with a Biotage Sfär Silica column, using a gradient of HPLC grade acetonitrile and water as mobile phase. A second round of purification was performed using an Agilent 1260 Infinity II HPLC coupled with an Agilent 1290 Infinity II Diode Array Detector. Separation was achieved using a reversed phase Agilent preparatory column (5 µm, 100Å C18, 21.2 x 150 mm) with a flow rate of 10 ml min^-1^. The mobile phase was a gradient mixture of acetonitrile and formic acid in water following the elution programme: 0 minutes, 5% B; 5 minutes, 5% B; 30 minutes 95% B, 35 minutes 95% B, 45 minutes 5% B (A, HPLC grade H_2_O containing 0.1% formic acid; B, HPLC grade acetonitrile containing 0.1% formic acid). Target compounds were isolated based on UV absorption, collected in glass vials prior to being pooled and concentrated *in vacuo*.

**NMR spectroscopy analysis of purified metabolites**

NMR spectroscopy was used to characterise purified metabolites. Dried compounds were dissolved in CDCl_3_ and characterised by NMR spectroscopy, which was conducted on an AV400 Bruker Avance III 400 MHz instrument. Chemical shifts were recorded in parts per million (ppm) and coupling constant (*J*) in Hz.

**Stable isotope feeding**

Starter cultures of *H. grisea* were set up using fungal agar plugs as an inoculum in 10 ml of YM glucose broth for ten days and grown in an orbital shaker at 180 rpm at 25°C. A Qiagen TissueRuptor II was used to homogenise the mycelial growth and 500 µl were used as an inoculum to set up fresh 20-ml YM glucose broth cultures in an orbital shaker at 180 rpm at 25°C. These were fed with overall 2.5 mM of either *L*-phenyl-^13^C_9_-alanine or standard *L*-phenylalanine over three days (day 3, 5 and 7 post inoculum) and cultured overall for two weeks. Metabolite extractions were performed twice using two volumes of ethyl acetate, followed by concentration of the crude extract *in vacuo*. Crude extracts were dissolved in methanol and analysed on LC-HRMS with a Dionex 3000RS UHPLC coupled to a Bruker UHR Q-TOF Maxis II mass spectrometer with an electrospray source. Separation was achieved using a reversed phase Agilent ZORBAX Eclipse Plus column (1.8 µm, 95Å C18, 2.1 x 100 mm). Sodium formate (10 mM) was used as the calibration standard, and a *m/z* scan range of 50–1500 was used. Auto MS/MS analysis was performed, in which the three most intense peaks were selected for MS^2^ data acquisition after each full scan. The mobile phase was a gradient mixture of acetonitrile and formic acid in water following the elution programme: 0 minutes, 5% B; 5 minutes, 5% B; 25 minutes 100% B, 30 minutes 100% B, 37 minutes 5% B (A, HPLC grade H_2_O containing 0.1% formic acid; B, HPLC grade acetonitrile containing 0.1% formic acid).

***H. grisea* genome sequencing, assembly and annotation**

High-molecular weight (HMW) genomic DNA from *H. grisea* was extracted using 100 mg of frozen mycelia obtained through cryogenic grinding. A GenElute^TM^ Plant Genomic DNA Miniprep Kit (Merck) was used, following the manufacturer’s instructions, including the RNase A optional treatment. HMW genomic DNA was concentrated using ethanol precipitation. Briefly, 1/10 volume of 3 M sodium acetate pH 5.2, was added, followed by three volumes of 100% ethanol. Samples were inverted, incubated at −20°C overnight, then spun at 13,000× g for 30 minutes at 4°C. Supernatant was decanted, the DNA pellet was air dried, and then resuspended in nuclease-free water. Quality and integrity of the HMW genomic DNA were assessed using a combination of agarose gel electrophoresis, NanoDrop ND1000 spectrophotometer (Thermo Fisher Scientific) and Qubit 4 fluorometer (dsDNA Broad range, Invitrogen) analysis. The HMW genomic DNA was used for both Illumina short-read sequencing (2x150 bp sequencing, 10M read pairs; Genewiz) and Nanopore long-read sequencing (performed in house). Library preparation for Nanopore sequencing was performed using the ligation sequencing kit (SQK-LSK109). The manufacturer’s protocol was followed with minor modifications introduced for both DNA repair and end-prep stages, by increasing the incubation time and temperature after washing with ethanol to 15 minutes at 37°C. To enrich DNA fragments of 3kb or longer, the Long Fragment Buffer was used for the DNA clean-up. An amount of 1 µg of the genomic DNA was used for the library preparation, resulting in a final amount of 350 ng of DNA at the end of the adapter ligation and clean-up step. The resulting DNA library containing the sequencing buffer and loading beads were loaded on the primed SpotON flow cell. Nanopore sequencing was performed on MinION (ONT) with a FLO-MIN-106 R9.4 flow-cell (ONT) and run for 18 hours. The raw data produced by nanopore sequencing were used to *de novo* assemble the genome and annotate it following the same pipeline used previously by us for fungal genomes.^1^

**Transcriptomics analysis**

Starter cultures of *H. grisea* were set up using fungal agar plugs as an inoculum in 10 ml of YM broth with different carbon sources in parallel (3 g l^-1^ yeast extract, 5 g l^-1^ peptone, 13 g l^-1^ glucose/mannitol/lactose/galactose/fructose, pH 6.2) and grown for ten days in an orbital shaker at 180 rpm at 25°C. A Qiagen TissueRuptor II was used to homogenise the mycelial growth and 500 µl were used as an inoculum to set up fresh 50-ml cultures of the same respective medium in an orbital shaker at 180 rpm at 25°C. These were grown for two weeks, after which metabolite extractions and analysis on LC-HRMS were performed as described above. For RNA sequencing *H. grisea* was grown on YM glucose broth, as well as on *H. grisea* grown on YM mannitol broth, as described above. The 50-ml cultures of each medium were grown for one and two weeks, each one in triplicate for both carbon sources, after which the mycelia were collected through centrifugation at 3,000 rpm for 10 minutes and ground to a fine powder with the aid of liquid nitrogen. One hundred mg of ground mycelia were used to purify RNA using the Plant/Fungi total RNA purification kit (Norgen Biotek) following the manufacturer’s instructions, including the optional on-column DNAase treatment with the RNase-Free DNase I kit (Norgen Biotek). RNA concentration and quality were assessed using a combination of agarose gel electrophoresis and NanoDrop ND1000 spectrophotometer (Thermo Fisher Scientific) analysis prior to messenger RNA enrichment, complementary DNA library preparation and Illumina NovaSeq 2x150 bp sequencing provided by Source Bioscience. The RNAseq reads were mapped against the *de novo* assembled *H. grisea* genome using Trinity^2^ and differential gene expression was determined. The fold change and log2 fold change were calculated based on the Fragments Per Kilobase of transcript per Million mapped reads (FPKM) of glucose/mannitol-grown *H. grisea* for each gene at a given time point of collection of the respective RNA (Supplementary Dataset 1). A log2 fold change value of 1.5 was used as threshold to determine a significant change in gene expression.

**Heterologous gene expression in *A. oryzae* NSAR1**

Plasmid pDA001 was constructed through Gibson Assembly by joining the backbone plasmid pTYGSarg^3^ and the intron-less sequences of *HGRIS_005317* and *HGRIS_000139*, which were amplified from cDNA of *H. grisea* with Q5 polymerase (NEB) using primer pairs FA111/FA229 and FA230/FA231 (Supplementary Table 7), respectively, according to the manufacturer’s instructions. Protoplast-mediated transformation of plasmid pDA001 into *A. oryzae* NSAR1 was performed as reported previously,^4^ and selecting transformant strains on CZDS agar plates selective for arginine (35 g l^-1^ Czapek-dox broth premix, 1 M sorbitol, 1.5 g l^-1^ methionine, 0.1 g l^-1^ adenine, 2 g l^-1^ ammonium sulphate, 20 g l^-1^ agar). The insertion of heterologous genes in *A. oryzae* DA1 was confirmed through PCR amplification of *HGRIS_005317* and *HGRIS_000139* with DreamTaq DNA Polymerase (Thermo Fisher Scientific) using primer pairs FA249/FA250 and FA251/FA252 (Supplementary Table 7), respectively, according to the manufacturer’s instructions.

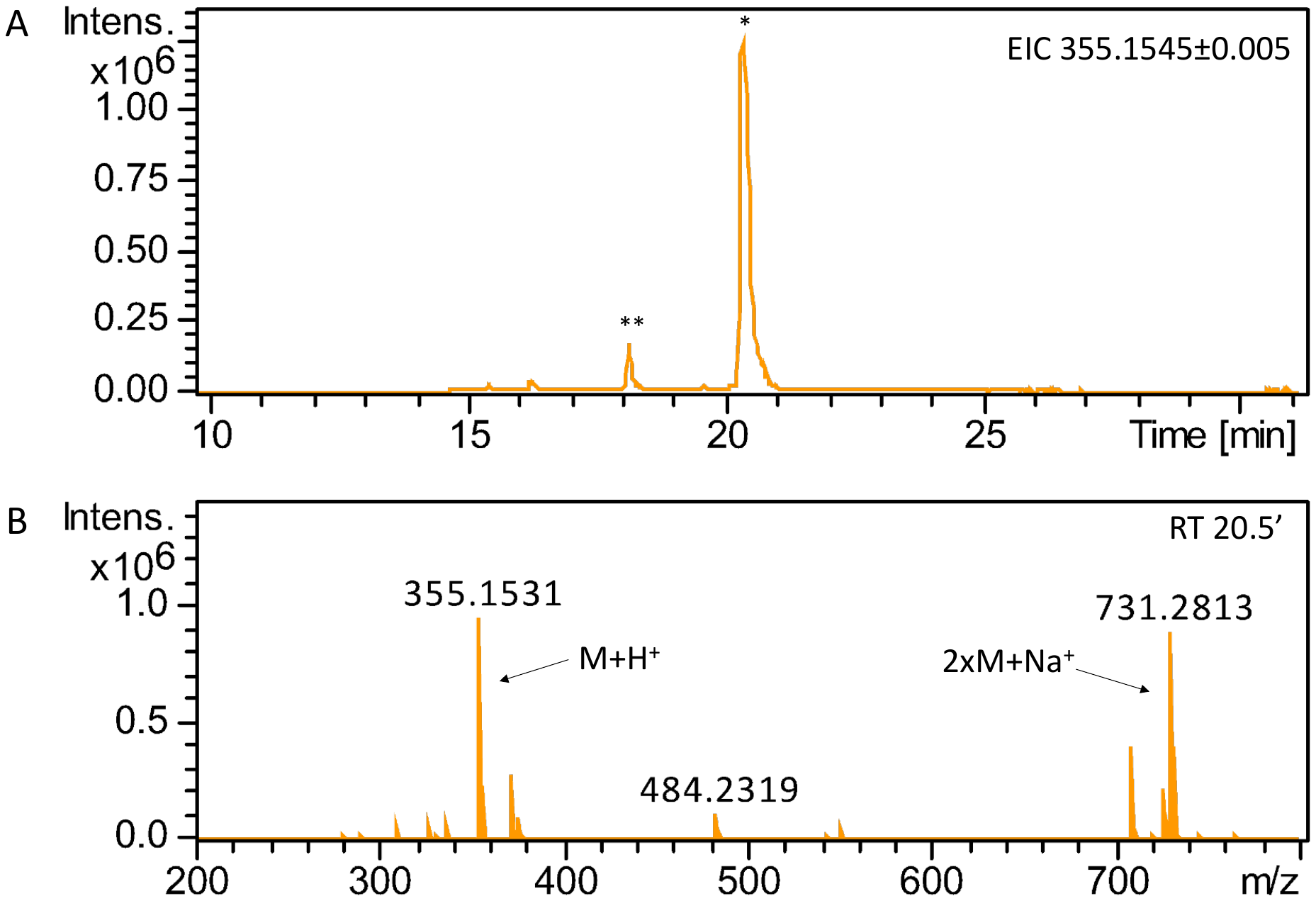

Supplementary Figure 1. LC-HRMS detection of pleurotin. **A**) Extracted ion chromatogram in positive mode at *m/z* = 355.1545±0.005 is shown, highlighting accumulation of pleurotin (**1**) in *H. grisea* crude extracts. One major peak (*) at retention time 20.5’ is detected, corresponding to **1**, based on NMR characterisation of the purified compound (Supplementary Figure 2); one additional peak (**) at retention time 18.2’ is detected, likely corresponding to nematoctone (**2**).^5^ **B**) Mass spectrum of pleurotin (M = C_21_H_22_O_5_), *m/z* expected for [M + H]^+^= 355.1545, detected at retention time 20.5’.

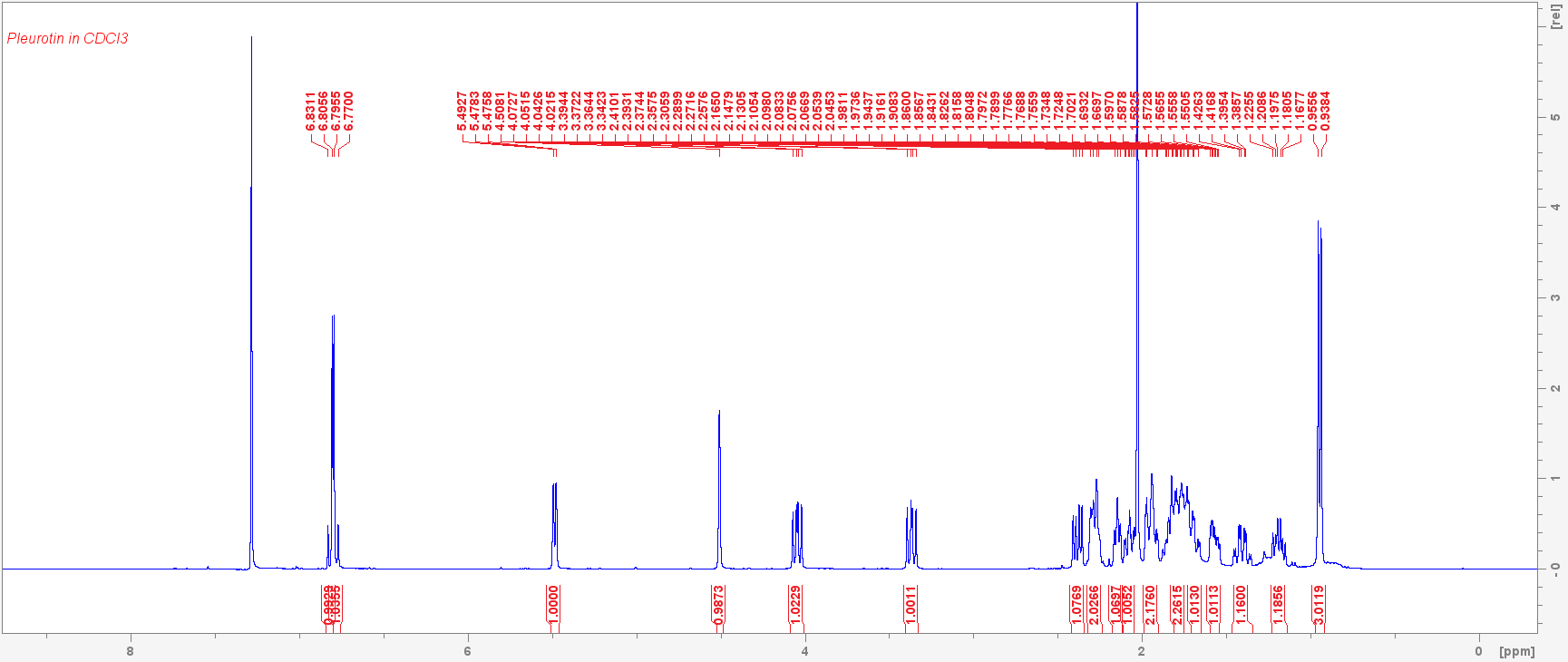

Supplementary Figure 2. ^1^H-NMR spectrum of 1 in CDCl_3_ (400 MHz). *δ* (ppm) 6.81 (d, *J* = 10.2 Hz, 1 H), 6.79 (d, *J* = 10.2 Hz, 1 H), 5.48 (d, *J* = 6.8 Hz, 1 H), 4.51 (s, 1 H), 4.05 (dd, *J* =8.5, 12.0 Hz, 1 H), 3.37 (dd, *J* = 8.9, 12.0 Hz, 1 H), 2.38 (dd, *J* =6.8, 14.3 Hz, 1 H), 2.31 – 2.23 (m, 2H), 2.15 (t, *J* = 6.9 Hz, 1 H), 2.11 – 2.05 (m, 1H), 1.99 – 1.90 (m, 2H), 1.82 – 1.74 (m, 2H), 1.68 (qd, *J* = 12.9, 3.9 Hz, 1H), 1.60 – 1.53 (m, 1H), 1.40 (qd, *J* = 12.3, 3.9 Hz, 1H), 1.19 (td, *J* = 11.6, 6.2 Hz, 1H), 0.95 (d, *J* = 6.9 Hz, 3 H) in agreement with published data for pleurotin.^6^

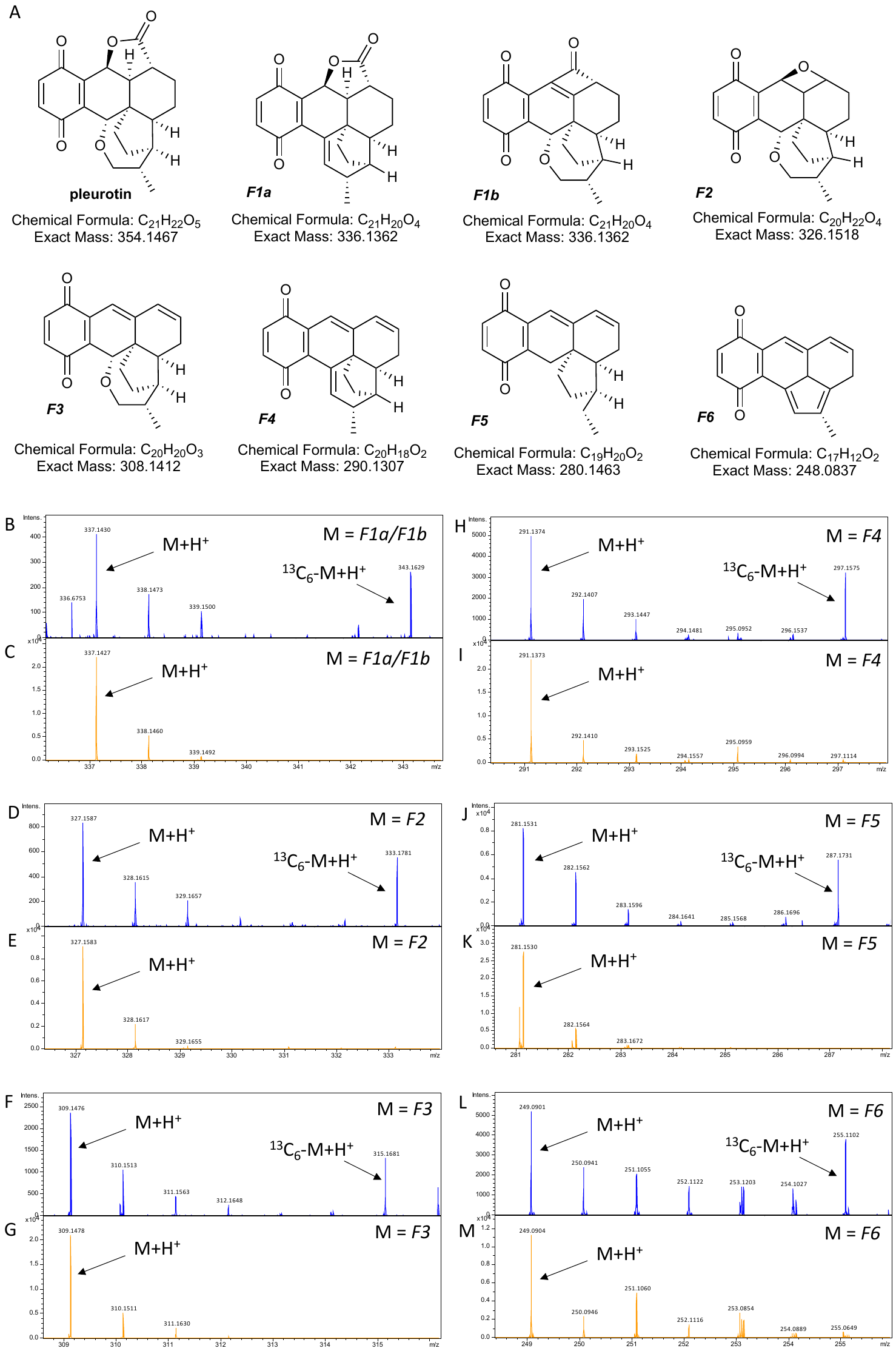

Supplementary Figure 3. MS^2^ spectra of *H. grisea* crude extracts grown in standard *L*-phenylalanine and in *L*-phenyl-^13^C_9_-alanine. Cultures grown in standard *L*-phenylalanine are shown by traces in orange (from parent ion *m/z* 355.1533 detected at retention time 20.5’) whereas cultures grown in *L*-phenyl-^13^C_9_-alanine are shown by traces in blue (from parent ion 361.1730 detected at retention time 20.5’) and show incorporation of six heavy carbons from *L*-phenyl-^13^C_9_-alanine in pleurotin. **A**) Structures of pleurotin and proposed fragments derived from it in MS^2^. **B**) MS^2^ spectrum showing species *F1a/F1b* and ^13^C_6_-*F1a/F1b*; *m/z* expected for *F1a/F1b* [M + H]^+^= 337.1439; *m/z* expected for ^13^C_6_-*F1a/F1b* [^13^C_6_-M + H]^+^= 343.1640. **C**) MS^2^ spectrum showing species *F1a/F1b*. **D**) MS^2^ spectrum showing species *F2* and ^13^C_6_-*F2*; *m/z* expected for *F2* [M + H]^+^= 327.1596; *m/z* expected for ^13^C_6_-*F2* [^13^C_6_-M + H]^+^= 333.1797. **E**) MS^2^ spectrum showing species *F2*. **F**) MS^2^ spectrum showing species *F3* and ^13^C_6_-*F3*; *m/z* expected for *F3* [M + H]^+^= 309.1490; *m/z* expected for ^13^C_6_-*F3* [^13^C_6_-M + H]^+^= 315.1691. **G**) MS^2^ spectrum showing species *F3*. **H**) MS^2^ spectrum showing species *F4* and ^13^C_6_-*F4*; *m/z* expected for *F4* [M + H]^+^= 291.1385; *m/z* expected for ^13^C_6_-*F4* [^13^C_6_-M + H]^+^= 297.1586. **I**) MS^2^ spectrum showing species *F4*. **J**) MS^2^ spectrum showing species *F5* and ^13^C_6_-*F5*; *m/z* expected for *F5* [M + H]^+^= 281.1541; *m/z* expected for ^13^C_6_-*F5* [^13^C_6_-M + H]^+^= 287.1742. **K**) MS^2^ spectrum showing species *F5*. **L**) MS^2^ spectrum showing species *F6* and ^13^C_6_-*F6*; *m/z* expected for *F6* [M + H]^+^= 249.0915; *m/z* expected for ^13^C_6_-*F6* [^13^C_6_-M + H]^+^= 255.1116. **M**) MS^2^ spectrum showing species *F6*.

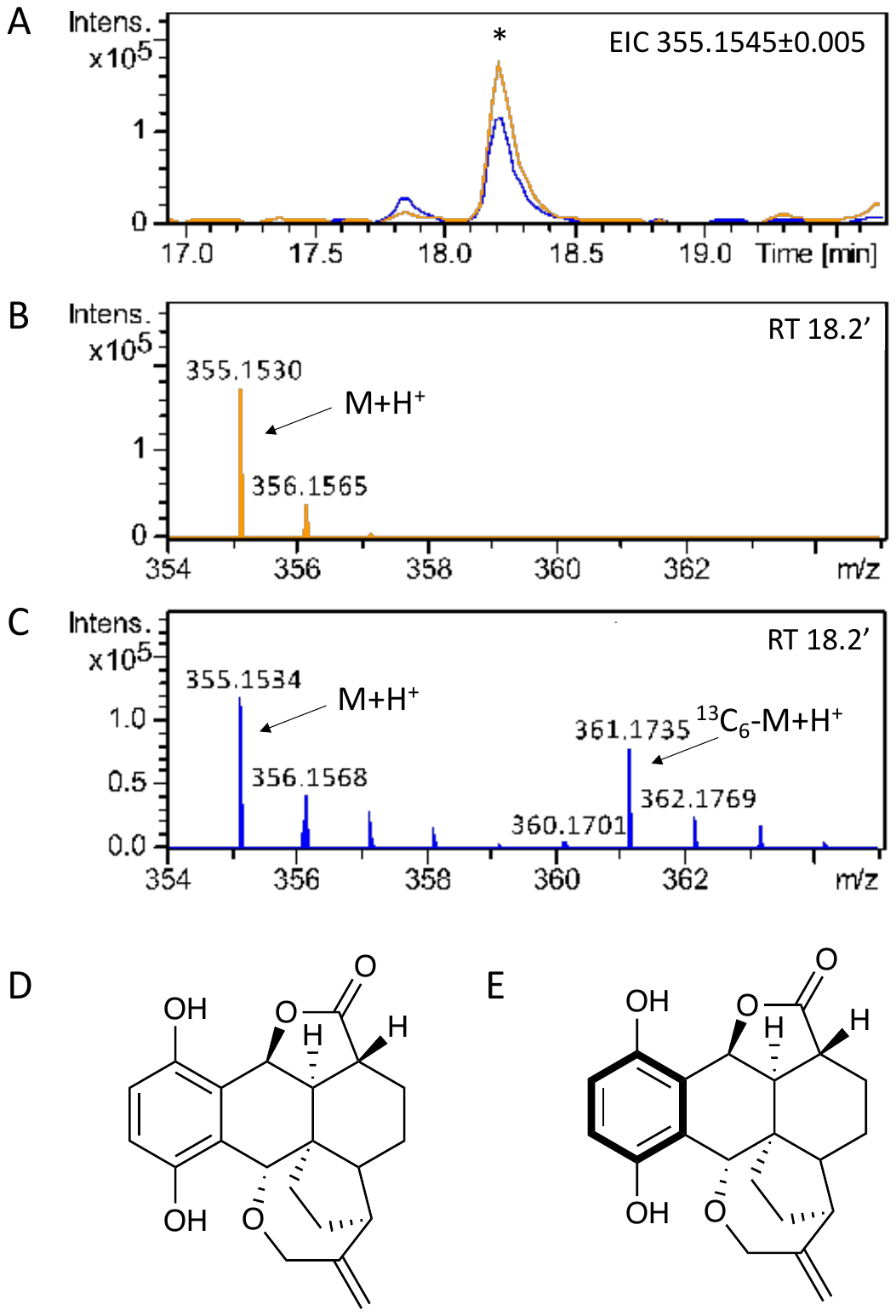

Supplementary Figure 4. LC-HRMS detection of nematoctone (2). **A**) Extracted ion chromatogram in positive mode at *m/z* = 355.1545±0.005 is shown, highlighting accumulation of nematoctone in *H. grisea* crude extracts grown in standard *L*-phenylalanine (trace in orange) and in *L*-phenyl-^13^C_9_-alanine (trace in blue). **B**) Mass spectrum of nematoctone (M= C_21_H_22_O_5_) detected at retention time 18.2’; *m/z* expected for [M + H]^+^= 355.1545. **C**) Mass spectrum of nematoctone and ^13^C_6_-nematoctone detected at retention time 18.2’; *m/z* expected for [^13^C_6_-M + H]^+^= 361.1746. **D**) nematoctone. E) ^13^C_6_-nematoctone.

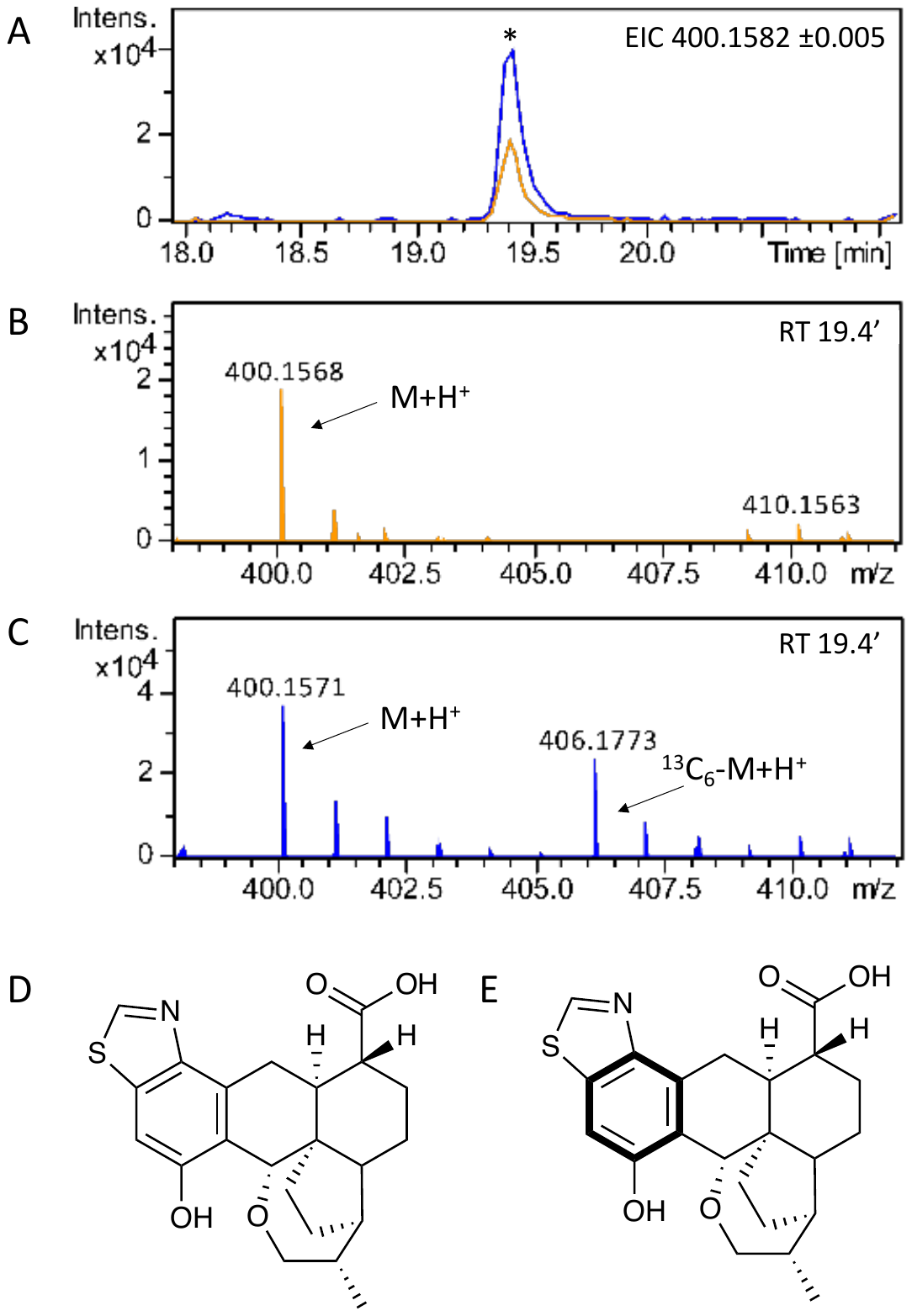

Supplementary Figure 5. LC-HRMS detection of pleurothiazole (4). **A**) Extracted ion chromatogram in positive mode at *m/z* = 400.1582 ±0.005 is shown, highlighting accumulation of pleurothiazole in *H. grisea* crude extracts grown in standard *L*-phenylalanine (trace in orange) and in *L*-phenyl-^13^C_9_-alanine (trace in blue). **B**) Mass spectrum of pleurothiazole (M= C_22_H_25_NO_4_S) detected at retention time 19.4’; *m/z* expected for [M + H]^+^= 400.1582. **C**) Mass spectrum of pleurothiazole and ^13^C_6_-pleurothiazole detected at retention time 19.4’; *m/z* expected for [^13^C_6_-M + H]^+^= 406.1783. **D**) pleurothiazole. E) ^13^C_6_-pleurothiazole.

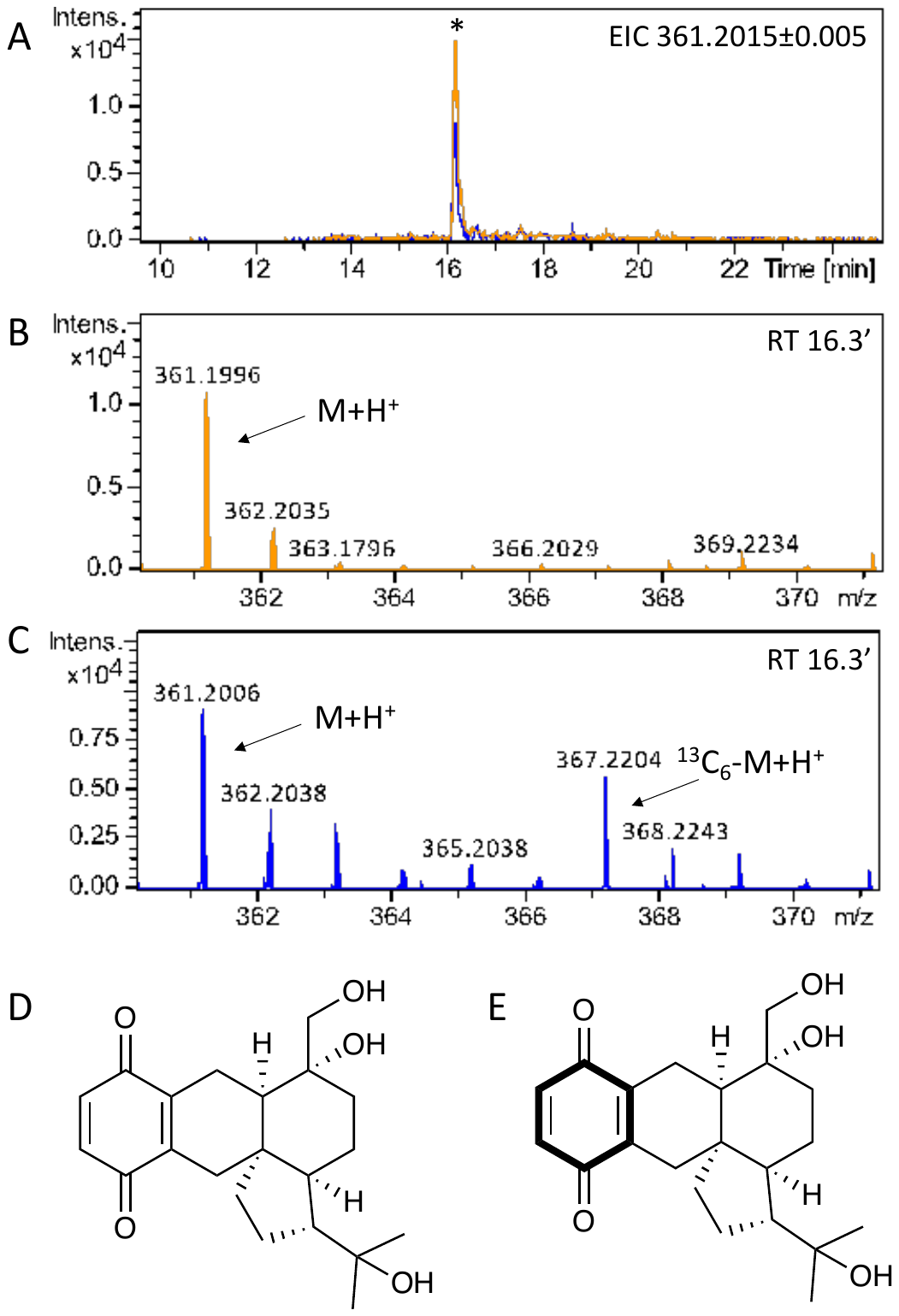

Supplementary Figure 6. LC-HRMS detection of 4-hydroxypleurogrisein (5). **A**) Extracted ion chromatogram in positive mode at *m/z* = 361.2015±0.005 is shown, highlighting accumulation of 4-hydroxypleurogrisein in *H. grisea* crude extracts grown in standard *L*-phenylalanine (trace in orange) and in *L*-phenyl-^13^C_9_-alanine (trace in blue). **B**) Mass spectrum of 4-hydroxypleurogrisein (M= C_21_H_28_O_5_) detected at retention time 16.3’; *m/z* expected for [M + H]^+^= 361.2015. **C**) Mass spectrum of 4-hydroxypleurogrisein and ^13^C_6_-4-hydroxypleurogrisein detected at retention time 16.3’; *m/z* expected for [^13^C_6_-M + H]^+^= 367.2216. **D**) 4-hydroxypleurogrisein. E) ^13^C_6_-4-hydroxypleurogrisein.

### Supplementary Table 1. Percentage of ^13^C incorporation from *L*-phenyl-^13^C_9_-alanine into pleurotin and three congeners.

| Compound | Calculated *m/z* | % ^13^C incorporation | Retention time |
| --- | --- | --- | --- |
| pleurotin | 355.1545 | 68 | 20.5’ |
| nematoctone* | 355.1545 | 67 | 18.2’ |
| pleurothiazole* | 400.1582 | 65 | 19.4’ |
| 4-hydroxypleurogrisein* | 361.2015 | 63 | 16.3’ |

* The identity of nematoctone, pleurothiazole and 4-hydroxypleurogrisein was deduced based on a match between the predicted molecular formulae from HRMS data and the known formulae of the compounds.^5,7^

### Supplementary Table 2. Assembly statistics of the genome of *H. grisea*.

| Parameter | Value |
| --- | --- |
| Number of contigs | 25 |
| Total contigs length | 38,874,855 |
| Mean contig size | 15,549,942 |
| Longest contig | 4,677,384 |
| Shortest contig | 19,611 |
| Contigs > 10K nt | 23 (100%) |
| Contigs > 100K nt | 17 (68%) |
| Contigs > 1M nt | 13 (52%) |
| N50 | 2,881,227 |
| L50 | 5 |
| N80 | 2,035,287 |
| L80 | 10 |
| Genes | 14,934 |
| Protein-coding genes | 14,602 |
| exons | 100,367 |
| tRNAs | 332 |

Supplementary Table 3. BUSCO analysis of *H. grisea* to assess the quality of genome scaffold and predicted proteome. The Agaricales_odb10 database was used as a reference.

| Parameter | Value |
| --- | --- |
| *BUSCO Genome Statistics* |  |
| Percentage BUSCO | 97.30% |
| Complete BUSCO’s | 3768 |
| Complete and single copy BUSCO’s | 3748 |
| Complete and duplicate BUSCO’s | 20 |
| Fragmented BUSCO’s | 3 |
| Missing BUSCO’s | 99 |
| Total BUSCO groups searched | 3870 |
| *BUSCO Predicted Proteome Statistics* |  |
| Percentage BUSCO | 94.30% |
| Complete BUSCO’s | 3648 |
| Complete and single copy BUSCO’s | 3343 |
| Complete and duplicate BUSCO’s | 305 |
| Fragmented BUSCO’s | 99 |
| Missing BUSCO’s | 123 |
| Total BUSCO groups searched | 3870 |

Supplementary Table 4. BlastP analysis of the *H. grisea* predicted proteome using Saccharomyces cerevisiae COQ2 as a query. The length of the *S. cerevisiae* COQ2 protein is 372 amino acids.

| Protein ID | Length (amino acids) | Identities | Positives | Gaps |
| --- | --- | --- | --- | --- |
| HGRIS_009700 | 379 | 124/277 (45%) | 171/277 (62%) | 5/277 (2%) |
| HGRIS_000873 | 327 | 97/293 (33%) | 152/293 (52%) | 17/293 (6%) |
| HGRIS_000139 | 309 | 100/301 (33%) | 150/301 (49%) | 5/301 (1%) |
| HGRIS_006930 | 247 | 89/250 (36%) | 124/250 (50%) | 24/250 (10%) |

Supplementary Table 5. BlastP analysis of the *H. grisea* predicted proteome using Boreostereum vibrans VibMO1 as a query. The length of the *B. vibrans* VibMO1 protein is 451 amino acids.

| Protein ID | Length (amino acids) | Identities | Positives | Gaps |
| --- | --- | --- | --- | --- |
| HGRIS_002929 | 468 | 186/442 (42%) | 250/442 (56%) | 25/442 (5%) |
| HGRIS_001008 | 459 | 122/428 (29%) | 197/428 (46%) | 16/428 (3%) |
| HGRIS_001007 | 479 | 121/427 (28%) | 190/427 (44%) | 18/427 (4%) |
| HGRIS_004802 | 609 | 118/431 (27%) | 203/431 (47%) | 24/431 (5%) |
| HGRIS_014061 | 470 | 122/433 (28%) | 199/433 (45%) | 25/433 (5%) |
| HGRIS_008360 | 426 | 120/403 (30%) | 192/403 (47%) | 31/403 (7%) |
| HGRIS_010717 | 457 | 130/464 (28%) | 213/464 (45%) | 60/464 (12%) |

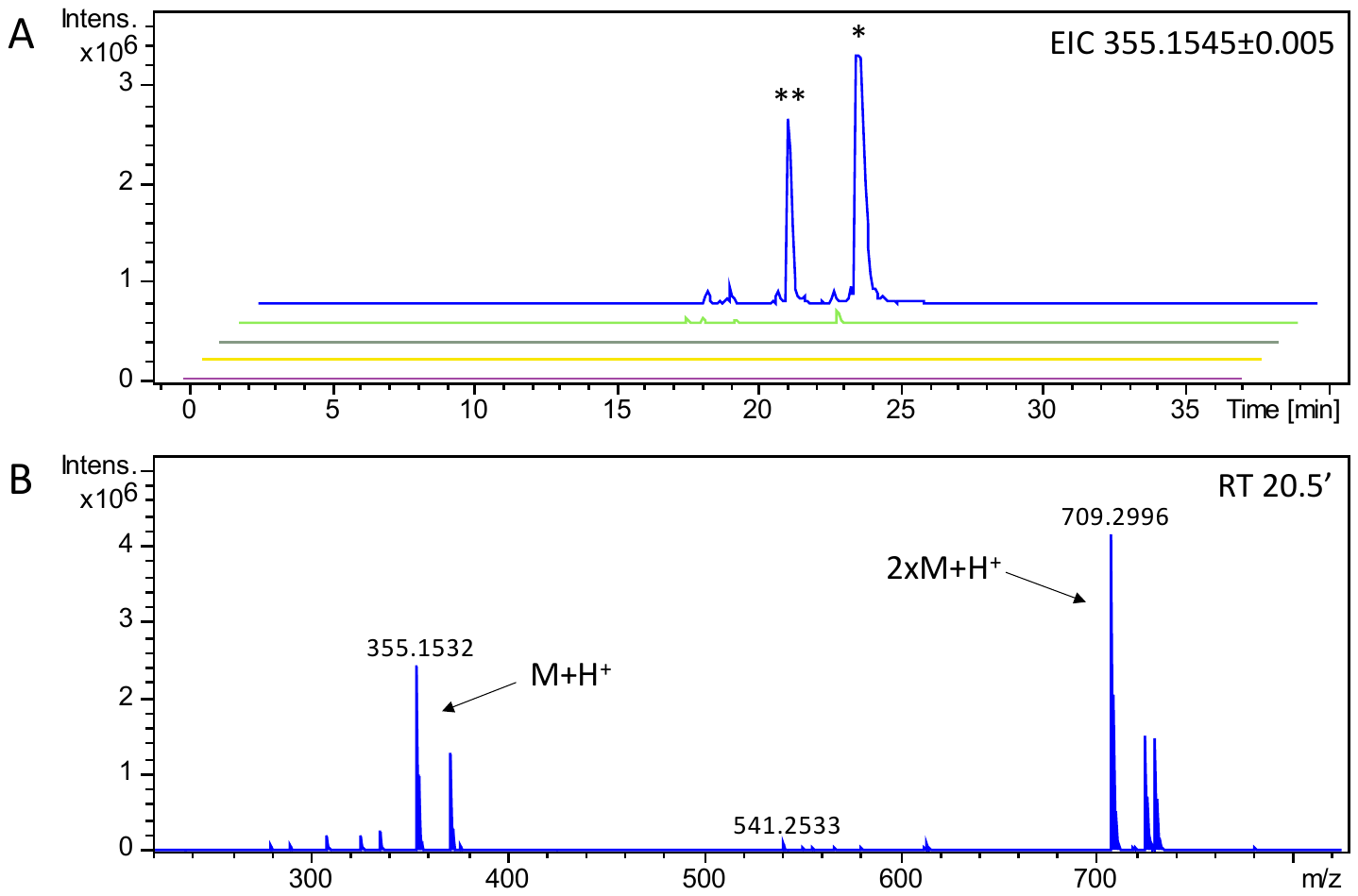

Supplementary Figure 7. LC-HRMS detection of pleurotin from *H. grisea* cultures grown in YM broth containing different carbon sources. **A**) Extracted ion chromatogram in positive mode at *m/z* 355.1545±0.005 for mycelia grown in mannitol (purple), lactose (yellow), galactose (grey), fructose (green), and glucose (blue) as the carbon source. One major peak (*) at retention time 20.5’ is seen, corresponding to pleurotin, based on NMR characterisation of the purified compound; one additional peak (**) at retention time 18.2’ is seen, likely corresponding to nematoctone.^5^ **B**) Mass spectrum of pleurotin (M= C_21_H_22_O_5_), *m/z* expected for [M + H]^+^= 355.1545, from mycelia grown in glucose as the carbon source, detected at retention time 20.5’.

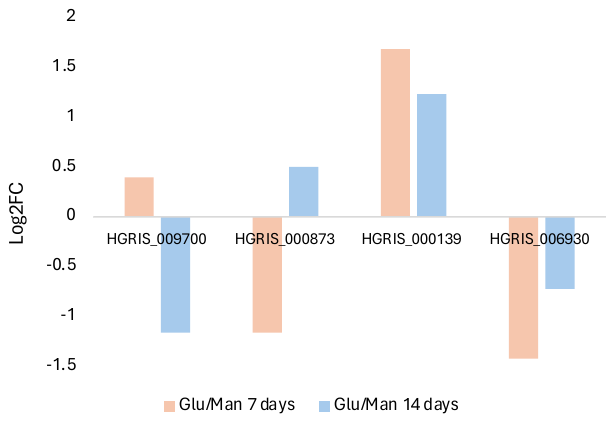

Supplementary Figure 8. Relative gene expression levels of *H. grisea* COQ2 homologues under pleurotin producing and non-producing fermentation conditions. Data at day 7 (in orange) and at day 14 (in blue) are shown. Log_2_FC= Log_2_ fold change of gene expression in glucose-grown fungi over mannitol-grown fungi.

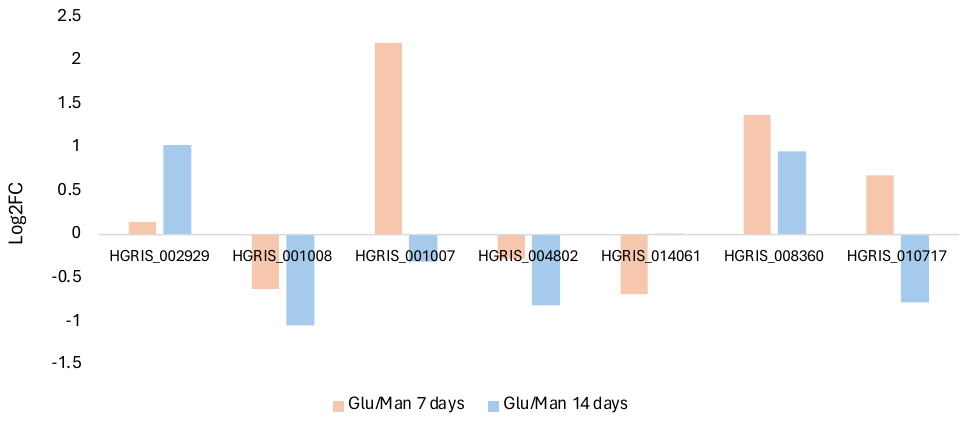

Supplementary Figure 9. Relative gene expression levels of *H. grisea* VibMO1 homologues under pleurotin producing and non-producing fermentation conditions. Data at day 7 (in orange) and at day 14 (in blue) are shown. Log_2_FC= Log_2_ fold change of gene expression in glucose-grown fungi over mannitol-grown fungi.

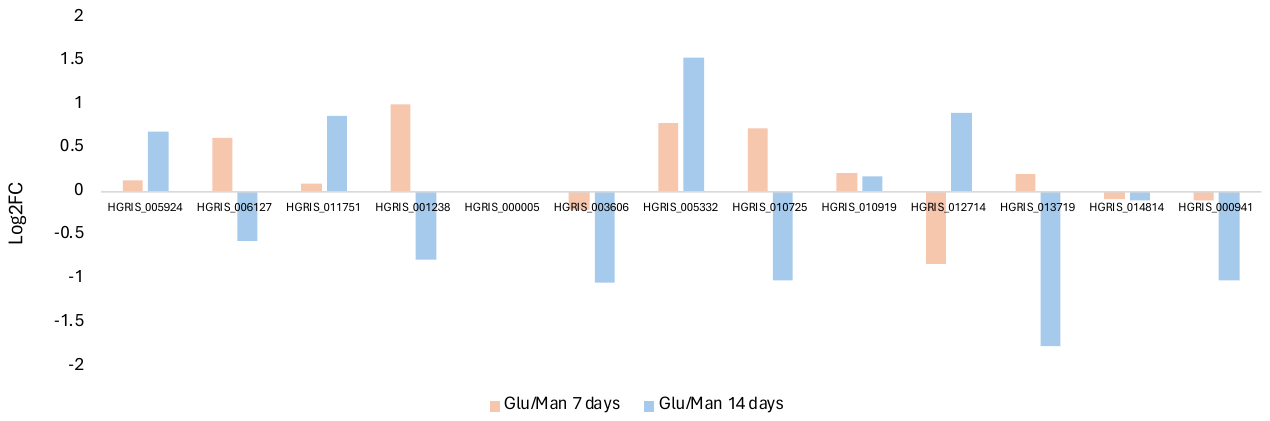

Supplementary Figure 10. Relative gene expression levels of *H. grisea* terpene synthases from FungiSMASH BGCs under pleurotin producing and non-producing fermentation conditions. Data at day 7 (in orange) and at day 14 (in blue) are shown. Log_2_FC= Log_2_ fold change of gene expression in glucose-grown fungi over mannitol-grown fungi.

### Supplementary Table 6. BlastP analysis of the translated genes from the BGC in region 8.1 from the genome of *H. grisea*.

| Protein ID | Length (amino acids) | Homologue (% identity/% similarity) | Organism (GeneBank ID) |
| --- | --- | --- | --- |
| HGRIS_005329 | 224 | Ferric/cupric reductase transmembrane component B (32/46) | *Aspergillus fumigatus* Af293 (Q4WR75.2) |
| HGRIS_005330 | 377 | Ferric/cupric reductase transmembrane component 7 (35/52) | *Saccharomyces cerevisiae* S288C (Q12333.2) |
| HGRIS_005331 | 608 | Ferric/cupric reductase transmembrane component 7 (31/48) | *Saccharomyces cerevisiae* YJM789 (A6ZN61.1) |
| HGRIS_005332 | 515 | Squalene synthase (69/80) | *Ganoderma lucidum* (A0SJQ5.1) |
| HGRIS_005333 | 523 | Acetylcholinesterase (34/47) | *Tetronarce californica* (P04058.2) |

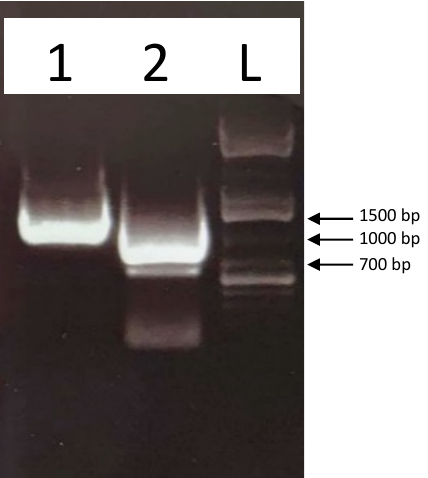

Supplementary Figure 11. PCR amplification of genes *HGRIS_005317* and *HGRIS_000139*. Primer pairs FA111/FA229 and FA230/FA231 were used to amplify *HGRIS_005317* and *HGRIS_000139*. Lane 1 contains amplicon *HGRIS_005317* (expected size 1,168 bp). Lane 2 contains amplicon *HGRIS_000139* (expected size 990 bp).

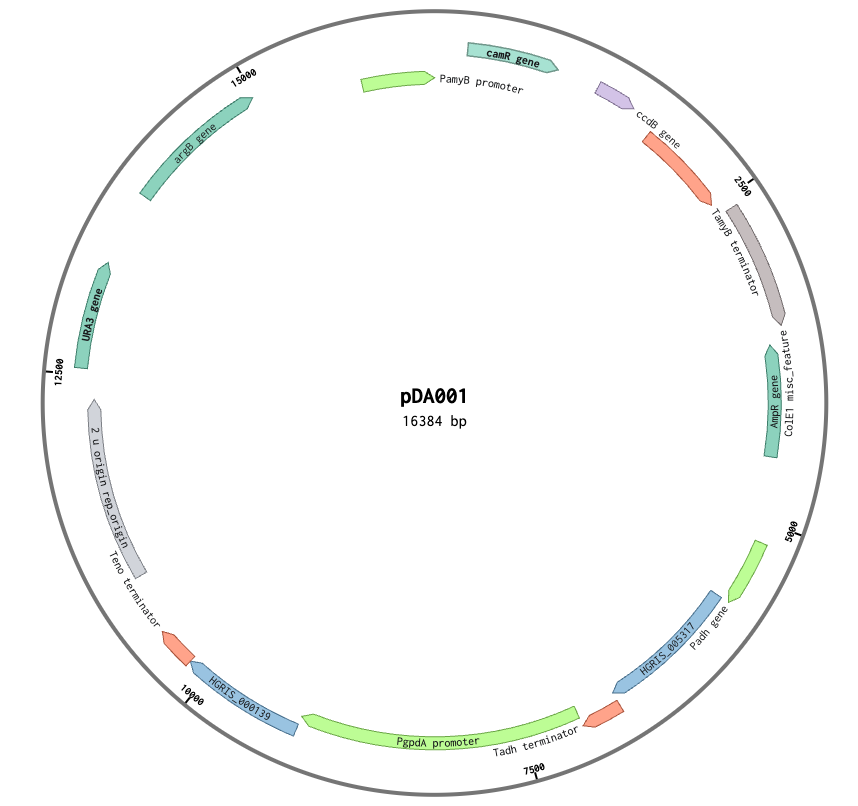

Supplementary Figure 12. Plasmid map of pDA001. The plasmid is based on pTYGSarg^3^ and includes *H. grisea* genes *HGRIS_005317* and *HGRIS_000139*.

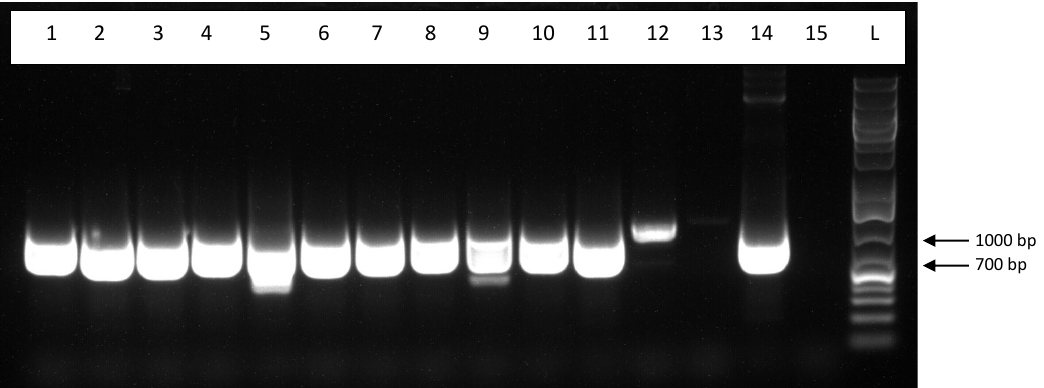

Supplementary Figure 13. PCR amplification screening of gene *HGRIS_005317*. Lanes 1-12 result from genomic DNA extracted from *A. oryzae* DA1 strains 1-12. Lane 13 results from genomic DNA extracted from recipient strain *A. oryzae* NSAR1. Lane 14 results from plasmid pDA001 (positive control). Lane 15 results from water (negative control). Expected amplicon size: 937 bp. The amplification was conducted using the primer pair FA251/FA252. Lane L contains 2 µL of GeneRuler 1kb plus DNA Ladder (Thermo Scientific).

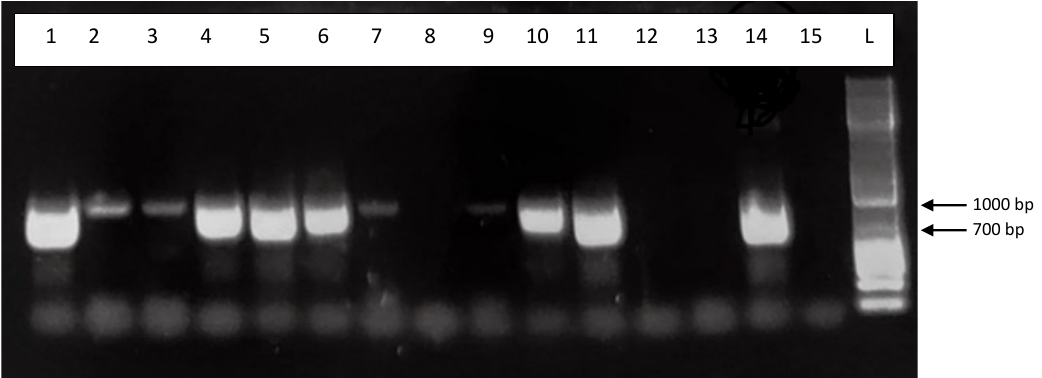

Supplementary Figure 14. PCR amplification screening of gene *HGRIS_000139*. Lanes 1-12 result from genomic DNA extracted from *A. oryzae* DA1 strains 1-12. Lane 13 results from genomic DNA extracted from recipient strain *A. oryzae* NSAR1. Lane 14 results from plasmid pDA001 (positive control). Lane 15 results from water (negative control). Expected amplicon size: 890 bp. The amplification was conducted using the primer pair FA249/FA250. Lane L contains 2 µL of GeneRuler 1kb plus DNA Ladder (Thermo Scientific).

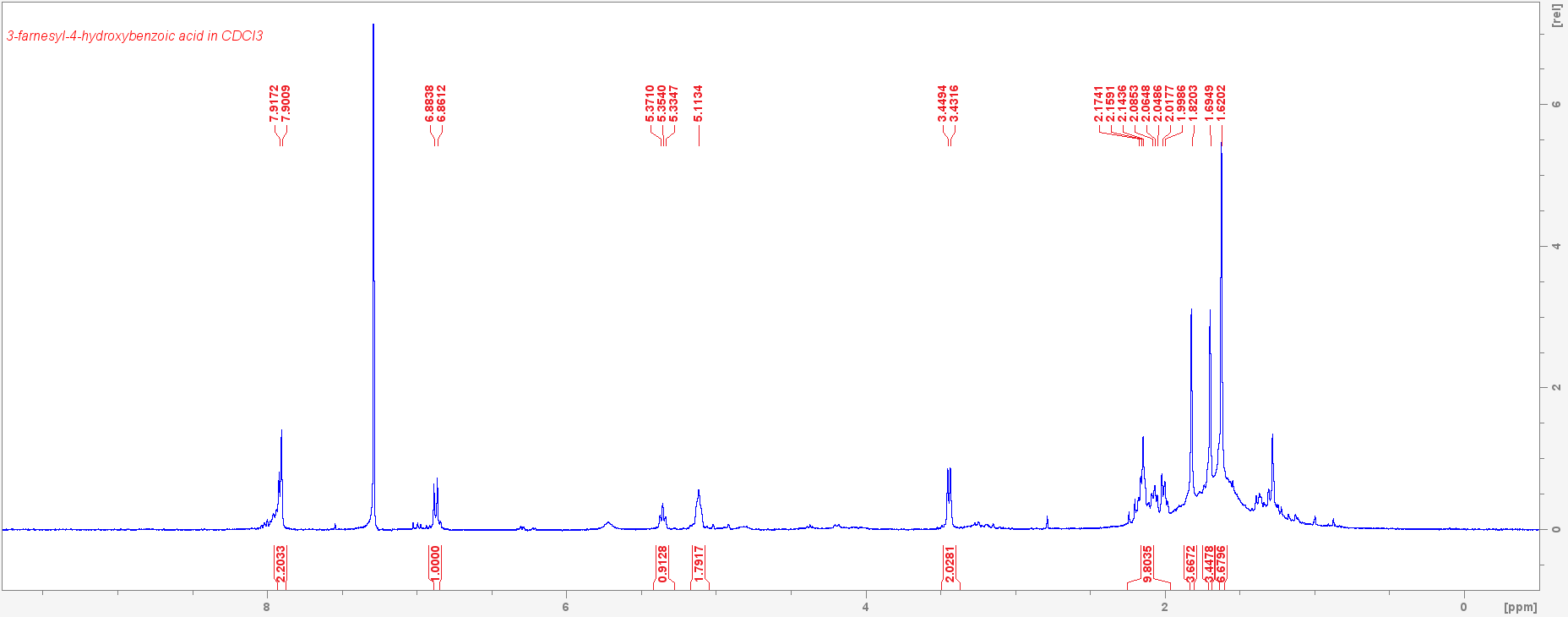

Supplementary Figure 15. ^1^H-NMR spectrum of 6 in CDCl_3_ (400 MHz). The signal observed at 7.26 ppm is from the hydrogen contained in CDCl_3_. *δ* (ppm) 7.90 (m, 2H), 6.88 (d, J= 9.0 Hz, 1H), 5.35 (t, J= 7.26 Hz, 1H), 5.11 (m, 2H), 3.44 (d, J= 7.1 Hz, 2H), 2.25–1.96 (m, 10H), 1.82 (s, 3H), 1.69 (s, 3H), 1.62 (s, 6H), in agreement with published data for 3-farnesyl-4-HBA.^8,9^

### Supporting Table 7. List of oligonucleotides and DNA sequences used in this work.

| Name | Sequence (5’ -> 3’) | Description |
| --- | --- | --- |
| FA111 | tttctttcaacacaagatcccaaagtcaaaATGGCTAGCACAAGCGCA | Amplification of *HGRIS_005317* for cloning of the intron-less gene into pTYGSarg^3^ |
| FA229 | tttcattctatgcgttatgaacatgttccctTCACTTCGACCTCTTGTAAATC |  |
| FA230 | aacagctaccccgcttgagcagacatcaccATGTCAGTTCTCAAGTCTCT | Amplification of *HGRIS_000139* for cloning of the intron-less gene into pTYGSarg^3^ |
| FA231 | ggttggctggtagacgtcatataatcatacTTACACGGCATGAGGAAGA |  |
| FA249 | AGTCTCTTCCCGCTTCTCTC | Amplification of *HGRIS_000139* for verification of plasmid insertion |
| FA250 | ACGCATTGACCAAAGGTAGT |  |
| FA251 | AGAACCTCGATTGGAATGTC | Amplification of *HGRIS_005317* for verification of plasmid insertion |
| FA252 | CTCGTGAACACCTGTCTCTT |  |
| HGRIS_005317 | ATGGCTAGCACAAGCGCAGACAAGAAGGCTGAGCGCCGTTCCCGATTCGAGGCTGCCTGGAATGTGATCCGCGACGAGCTCGTTGAACACCTCAAGGGTGAGGGTATGCCGGCCGATGCGCAAGAGTGGTTTAAGAAGAACCTCGATTGGAATGTCCCAGGCGGCAAGCTCAACCGCGGCATGTCCGTCGTCGACACTGTCGAGATCCTCAAAGCCCGTTCGCTGACCGAAGAGGAATACTTCAAGGCCGCTATTCTTGGCTGGTGTGTGGAGCTGCTCCAAGCGTACTTCCTCGTAGCCGATGACATGATGGACCAATCGATCACGCGCCGCGGCCAGCCTTGCTGGTACCGCGTCGAAAGCGTTGGCAATATTGCGATCAACGACTCGTTCATGCTCGAAGCTGCCATCTACTGGCTCCTCAAAAAGCATTTCCGCACCGAGCCATGCTACGTTGACCTCCTTGAGCTCTTCCACGAGACAACATATCAAACGGAGATGGGCCAGCTGATCGACTTGATCACCGCGCCAGAAGACTCTGTTGACCTCTCCAAATTCTCGCTCGAGAAGCACCGCCTCATCGTCGTCTACAAGACCGCCTTCTACTCCTTCTATCTTCCCGTAGCGCTCGCGATGTACCTCGCCGGCGTGCCGCCGTCCTACGTCCCTGCCGCTGCTCCTCATGCCGCTGCCGTGTCACCGTACAAGATCGCGCTCGACATCCTCCTGCCTCTGGGCGAGTACTTCCAGGTCCAAGACGACTTCCTCGACTACTCCCAGCCACCGGAGCTCCTTGGCAAGATTGGCACAGACATCATCGACAACAAGTGCTCGTGGTGCATCAACACCGCGCTTGCCAAGGTCACGCCTGAGCAGCGCGCGGTGCTCGACGCGTCATACGGGCGCAAGGACGCAGAGAAGGAGCAGCAGGTGAAGAAGATTTTCAAGGAGGTTGGCGTTGAGAAGGCGTATGAGGAGTATGAGGAGGCGGTTGTTGGCAAGATTCGCAAGATGATCGAGGCGGTTCCTGAGGGCACGAAGGAGGGCGAGTTGAAGAGACAGGTGTTCACGAGCTTTTTGGAGAAGATTTACAAGAGGTCGAAGTGA | Gene sequence amplified from cDNA of *H. grisea* and included into plasmid pDA001 |
| HGRIS_000139 | ATGTCAGTTCTCAAGTCTCTTCCCGCCTCTCTCCAGCCTTGGGTCGATTTGATGCGCCTCAGCAGGTATGCGGGGACAATGCTGATTTTTTGGCCATATGCCTGGGGAGCAACAATGGCTGCCCGGACCTCTCTACTTCCCGTGCCTCGGTTCTTTGAACTCATCGCCTATGGGCTCATTGGGGCTTCACTCGCGCATAGCGCTGGATGTGTCTGGAATGATATTCTTGACCGAAACTTTGACCGGAAAGTTGCGCGAACGAAGTCCCGCCCCATTGCTGCGGGGCTTATTTCTGTTCCAAAGGCTCTAGTCTTCTTGGCTGCCCACTATGCCTGCATGTTCTACATGATCTGGACAGTCAATCCTCTCGCCTGGAAGGTCGCCCTTACCACAATGTTCCCTCTGGCTGGTGCATACCCATTTATGAAGCGCATCACAAACTGGCCACAAGCCTGGCTAGGAATCTCTATGAACACTGGTATCCTCATGGCGTGGGCATTTATTGCTGGCGACGTCCCAAAGAGCTCTTGGGTTTTGCTTGGTGGAGCATGGGCATGGACAATTTGGTATGATACCATCTATGCCTGCCAAGACAAGAAGGACGACAAAAAGGCCGGCGTGAAGTCGACCGCACTCCTCTTCGGCGACAACATCAAAGCTGTCCTCGGAACGTTCGCTGCCCTTGTGGTTGGAGCGCTCTATTACGCTGGCAAGCTGAACGACCAGCAGTTGCCTTACTTCCTCATCAGTGTCCTTGGCGGAGCAGTCCACATGGGCGTCGAGCTGTATACTGTTGACACTGATTCGCCCAAGAGCTGTTGGGCTGCTTTCCATCGCGCAGGTTTCCACCTTGGAGCGTTGATTTGGACCGGCTTCTTGGCTGACTACCTTTGGTCAATGCGTGGAACCATTCTTCCTCATGCCGTGTAA | Gene sequence amplified from cDNA of *H. grisea* and included into plasmid pDA001 |
